# A stable 15-member bacterial SynCom promotes *Brachypodium* growth under drought stress

**DOI:** 10.1101/2024.09.10.612297

**Authors:** Archana Yadav, Mingfei Chen, Shwetha M. Acharya, Yuguo Yang, Tiffany Z. Zhao, Romy Chakraborty

**Affiliations:** Department of Ecology, Earth & Environmental Sciences Area, Lawrence Berkeley National Laboratory, Berkeley, California, 94720, USA

**Keywords:** Microbiome, Rhizosphere, *Brachypodium*, Drought, SynCom

## Abstract

Rhizosphere microbiomes are known to drive soil nutrient cycling and influence plant fitness during adverse environmental conditions. Field-derived robust Synthetic Communities (SynComs) of microbes that mimic the diversity of rhizosphere microbiomes can greatly advance a deeper understanding of such processes. However, assembling stable, genetically tractable, reproducible, and scalable SynComs remains challenging. Here, we present a systematic approach using a combination of network analysis and cultivation-guided methods to construct a 15-member SynCom from the rhizobiome of *Brachypodium distachyon*. This SynCom incorporates diverse strains from five bacterial phyla and demonstrates strong stability both *in vitro* and *in planta*. Genomic analysis of the individual strains revealed that they encode multiple plant growth-promoting traits, some of which were validated by laboratory phenotypic assays. Additionally, most strains encoded genes both for the synthesis of osmoprotectants (trehalose and betaine) and Na^+^/K^+^ transporters. These traits likely enabled the resilience of *Brachypodium* to drought stress where plants amended with SynCom recovered better than without. We further observed preferential colonization of SynCom strains around root tips under stress, likely due to active interactions between plant root metabolites and bacteria. Our results represent significant progress towards building and testing stable model SynComs for a better understanding of plant-microbe interactions.

## Introduction

It is widely acknowledged that root-associated microorganisms play a crucial role in plant health and productivity. Root microbiomes aid plants by promoting nutrient acquisition (Mendes et al., 2013), producing plant growth hormones (Dodd et al., 2010; Eichmann et al., 2021; Joshi et al., 2021), and conferring resilience under abiotic stresses such as drought and increase in salinity (Lemanceau et al., 2017; Mayak et al., 2004; Schmitz et al., 2022; Yang et al., 2009). Plants recruit these root microbiomes primarily through the secretion of root exudates, which contains compounds such as organic acids, amino acids, nucleotides, vitamins, and fatty acids (Berendsen et al., 2018; Hartmann et al., 2008; Kawasaki et al., 2016; Mendes et al., 2013; Zhalnina et al., 2018).

Recent advances in sequencing technologies have provided valuable insights into the structure and function of root microbiomes across diverse plant species (Bulgarelli et al., 2015; Coleman, 2016; Liu et al., 2019, 2021). These studies have demonstrated that certain microbial groups are of more critical functional importance than others (Bergelson et al., 2019; Bulgarelli et al., 2012; Gkarmiri et al., 2017; Kwak et al., 2018). However, identifying and selecting beneficial root microbes for crop improvement remains challenging due to an incomplete understanding of complex interactions between microbes and their host plants. One promising approach is constructing simplified synthetic communities (SynComs) with defined interactions derived from root microbiomes (Gupta et al., 2021; Shayanthan et al., 2022). This method allows researchers to gain a deeper understanding of how microbes cooperate, compete, and influence each other and the plant host in the rhizosphere ecosystem (de Souza et al., 2020). SynComs are effective tools for unraveling intricate interactions to generate defensible hypotheses about microbial communities during plant growth and health under myriad environmental conditions. Generally, SynComs are designed either based on 1) a “Reductionist approach” that employs cultivation-based techniques to assemble SynComs (Liu et al., 2019; Rodríguez Amor, Dal Bello, 2019; Vorholt et al., 2017; Xia et al., 2020) or 2) a “Holistic approach” that focuses on assembling SynComs based on key microbial interactions in natural environment observed through culture-independent techniques like high-throughput sequencing technology (Gonçalves et al., 2023; Grosskopf, Soyer, 2014; Raes et al., 2011; Zegeye et al., 2019). However, both approaches have limitations. The reductionist approach often fails to recover key plant-associated microbes that are typically resistant to cultivation, or it may assemble a consortia of isolates that do not originate from the rhizosphere microbiome, and therefore have genotypic and functional differences compared to microbes naturally present in rhizosphere soil (Liu et al., 2019). On the other hand, the holistic approach which relies heavily on genome sequences, fails to distinguish between live and dead microbial cells leading to incorrect estimates of their diversity, and assumes that the presence of genomic signatures translates to functional activity (Gupta et al., 2021; Gutleben et al., 2018; Muller et al., 2013). Due to these inherent limitations, neither reductionist nor holistic approach alone is optimal. Therefore, a better approach to constructing SynCom that can encompass crucial microbe-microbe and microbe-host plant interactions to mimic the natural environment is warranted (Chodkowski, Shade, 2017; McCarty, Ledesma-Amaro, 2019).

Further, for broader utility, it is desirable that a SynCom is stable (with robust colonization and prevalence throughout plant development), effective (able to confer beneficial traits to plants), and reproducible (yield consistent results both in the laboratory experiments and during field studies). However, the successful assembly of such SynCom has been limited to a handful of prior studies (McCarty, Ledesma-Amaro, 2019; Shayanthan et al., 2022), with many SynComs failing to demonstrate comparable efficacy in the laboratory experiments versus field experiments in higher plants (Herrera Paredes et al., 2018; Nadeem et al., 2014; Zimmer et al., 2016). Thus, developing effective and stable SynComs has become a prominent area of interest in contemporary agriculture research, offering a potential natural, sustainable alternative to over-reliance on chemical fertilizers and improving crop productivity (Santhanam et al., 2015; Shayanthan et al., 2022; Wang et al., 2021).

In this study, we developed a stable and effective SynCom from a stable microbial community enriched from the root microbiome of *Brachypodium distachyon,* a well-characterized model grass species. We employed a combinatorial approach encompassing culture-independent root microbial abundance analysis and network analysis to identify critical microbe-microbe interactions, with culture-dependent isolation of key microbial strains, to construct a 15-member SynCom. We characterized the genomic and phenotypic traits of individual strains of SynCom and ascertained the SynCom’s stability in both liquid cultures as well as when grown with the plants. To further explore the SynCom’s stability and effectiveness in supporting plant growth under adverse environmental conditions, we amended *Brachypodium* seedlings with the SynCom before subjecting plants to salinity and drought stresses. Using 16S amplicon sequencing, whole genome sequencing, assays to assess plant-growth promoting traits and plant phenotyping, we discerned correlations between plant phenotype, SynCom abundance, and the beneficial effects of the SynCom on the plant growth under salinity and drought stress. Lastly, we investigated the spatial distribution of SynCom strains on roots to understand if these strains selectively colonize specific root niches.

## Materials and Methods

### Microbial isolation and identification

The 15-member SynCom was created using results from two interconnected experiments. In the first study (Acharya et al., 2023b), the microbe recruitment along the root surface was studied in young *Brachypodium* plants grown in natural soil. In the subsequent study (Chen et al., 2024), over 750 stable reduced community consortia (RCC) were enriched by growing root-attached microbes from the previous study over either 3 or 7 days in media with carbon substrates generally present in *Brachypodium* root exudates. In this study, based on high species diversity as denoted by the richness and Shannon diversity index, enrichments grown on two media i.e., 0.1X R2A (BD Diagnostics), and RCH2 minimal media (Chakraborty et al., 2017) supplemented with carbon sources (either Glutamine or mixed carbon), were used to isolate colonies on the corresponding agar media plates by incubating the plates at 30°C in the dark for 7 days. Morphologically distinct colonies were picked and streaked for further purification. For species identification of the isolates, genomic DNA extracted using a PureLink Genomic DNA Mini Kit (Invitrogen, United States), was amplified using the universal 16S rRNA eubacterial primer pairs i.e., 8F/27F and 1492R. DNA sequencing was carried out at the UC Berkeley DNA Sequencing Facility. Geneious Prime v2020.2.5 was used to process the 16S reads and the resulting consensus sequences were taxonomically classified using the SILVA database (Quast et al., 2013).

### Genomic analysis of individual isolates

The whole genome sequencing was performed at Novogene (Illumina Novaseq 6000 platform). Genome assembly and annotation were performed using the KBase platform (Arkin et al., 2018). Raw reads quality was assessed with FastQC v0.11.9, followed by trimming using Trimmomatic v0.36 (Bolger et al., 2014). The reads were then assembled using Spades v3.15.3 (Prjibelski et al., 2020) and the genome quality was evaluated using CheckM v1.0.18 (Parks et al., 2015). DRAM v0.1.2 (Shaffer et al., 2020) and Blastkoala (Kanehisa et al., 2016) were used to assign KEGG orthology (KO) numbers and functional annotation to the protein-coding genes and KEGG mapper (Kanehisa, Sato, 2020) was used to visualize metabolic pathways. Annotations from the metabolic assembly output files obtained from DRAM and KEGG were used to search for specific properties or genes such as PGP traits, transporters, and osmoprotectant-related genes. For the taxonomic classification, a concatenated alignment of 120 single-copy marker proteins was created using the GTDB-tk workflow (Chaumeil et al., 2022; Parks et al., 2022). Subsequently, 4-5 representative sequences closest to the SynCom strains were selected and aligned using clustalo (Sievers et al., 2011). This was followed by the tree construction with FastTree (Price et al., 2009) and visualization using iTOL (Letunic, Bork, 2007).

### Testing stability of SynCom members

We evaluated *in vitro* stability of the 15-member SynCom by tracking changes in the relative abundances of the strains over a three-week period. For this, we cultured 5 ml of the inoculum, containing equal cell numbers (4x10^7^ cells) of each SynCom member, in 45 ml of 0.2X MS (Murashige and Skoog) media (M0404, Sigma Aldrich, USA). The mixture was incubated in a plant growth chamber with conditions set at 16-hour light, 24℃ temperature, and 50% humidity. At the end of each week, 5 ml of the culture was transferred to 45 ml of fresh 0.2X MS media, and 5 ml of the culture was sampled for 16S rRNA community analysis.

In addition, to assess the resilience of the overall SynCom growth to the loss of individual members, we conducted a leave-one-out growth curve experiment. For this, individual SynCom members were first cultured in 0.1X R2A liquid medium until they reached the active growing stage. 15-member and 14-member consortia were created by mixing the liquid cultures of the individual members at a 1:1 (v:v) ratio. Next, 50 µL of each mixed consortium was inoculated into 450 µL of 1/10 R2A liquid medium in 48-well plates (n = 3). The optical density (OD) at 600 nm was recorded every hour using a BioTek Epoch 2 Microplate Spectrophotometer (Agilent Technologies, CA, USA) using Gen 6 1.04 software (Agilent Technologies, CA, USA).

### Seed germination and preparation of inoculum for in planta experiments

Approximately 150 *Brachypodium distachyon* (Bd 21-3) seeds were dehusked and sterilized by immersing in 70% ethanol for 30 seconds, 50% bleach for 5 minutes, and then washed five times with autoclaved MQ water. The sterilized seeds were stored in sterile water at 4℃ in the dark for one week. Subsequently, the seeds were germinated on semisolid plates containing 0.5X Murashige and Skoog Basal (MS) Medium (Sigma-Aldrich, USA) with 0.4% w/v phytagel and placed in a plant growth chamber (Percival Scientific AR-41L3, USA) set to 16 hours of light, 24℃ temperature, and 50% humidity for 3-5 days. For potting, sterilized plastic pots (3.9”D x 3.94” W x 3.15” H, Manufacturer: MiMiLai) lined with coffee filters and filled with autoclaved calcined clay (PROFILE Products LLC, USA) were used. The clay was saturated with 0.2X MS media before transplanting one seedling per pot. The weight of each pot, once filled with clay and plants and watered to clay saturation, was recorded as the saturation weight.

Individual SynCom isolates were cultivated in R2A media for 3-5 days until they reached log phase. After cultivation, the isolates were checked for purity, washed three times, and resuspended in a 30 mM phosphate buffer. For each isolate suspension, cells were stained with the nucleic acid stain SYBR green and counted using a flow cytometer (AttuneNxT®, ThermoFisher Scientific) following the manufacturer’s instructions. Suspensions for individual isolates were adjusted to equal cell numbers, specifically 4.1 x 10^7^ cells, for the final inoculum used in subsequent *in planta* experiments.

### Assessing the performance of SynCom under drought and salinity stress

This experiment comprised three conditions: Drought, Rewatered drought, and Salinity, to reflect natural environmental stresses. Each condition included two experimental sets; 1) SynCom-amended plants where 1 ml of 30 mM phosphate buffer containing equal cell numbers of each SynCom strain were inoculated at the base of plant shoot 3 days before inducing stress conditions, and 2) unamended plants where 1 ml of 30 mM phosphate buffer was inoculated at the base of plant shoot three days before inducing stress conditions. Each experimental set contained 7 plant replicates. After transplanting seedlings into pots, they were watered with 0.2X MS media to maintain 80% saturation weight for 4 days, after which they were inoculated with either SynCom or buffer, as explained earlier.

In addition to the three stress conditions, there was a control set with both SynCom-amended and unamended plants, maintained at 80% saturation weight by watering with 0.2X MS media. Stress induction began 3 days after SynCom inoculation. The ‘Drought’ condition involved maintaining plants at 40% saturation weight by watering with 0.2X MS media until they were harvested 21 days after stress induction. The ‘Rewatered drought’ condition involved 14 days of drought treatment, followed by restoring saturation weight to 80% for the last week before harvest. The ‘Salinity’ condition involved watering the plants at 80% saturation weight with 0.2X MS media containing 60 mM NaCl. After 3 weeks of stress induction, plants were harvested by gently removing them from the pots for phenotypic measurements, including shoot wet weight, root length, leaf count, and shoot length (longest leaf). In addition, root tip and root base samples were aseptically harvested as 2 cm cuttings and suspended in 5 ml of 5 mM sodium pyrophosphate + 30 mM phosphate buffer. The sample was sonicated for 10 minutes and subsequently left on a benchtop for 5 minutes to allow soil particles to settle down. The supernatant was pipetted out and pelleted to collect microbial cells. DNA extracted from these samples were sent to Novogene, USA, for amplicon sequencing. Quantitative Insights Into Microbial Ecology (QIIME2) (Bolyen et al., 2019) was used to process 16S rRNA amplicon data. Within QIIME2, DADA2 was employed for quality filtering, chimera checking, and paired-end read joining. The resulting sequences were trimmed to obtain V4 region (515F-806R), and taxonomic classification was performed using the SILVA database (Quast et al., 2013).

### Assay for plant growth promoting (PGP) traits in the SynCom strains

#### Biofilm formation assay

The crystal violet assay for quantifying biofilm formation was adapted from Haney et al. (Haney et al., 2021). Briefly, each isolate was grown in R2A media, washed, and resuspended in a 30 mM phosphate buffer to a final OD_600_ of 0.2. The cultures were inoculated into 96-well microtiter plates containing 180 µL of 0.2X MS media, at a 1:10 (v/v) ratio to achieve a final volume of 200 µL (initial OD_600_ of 0.02). The plates were incubated statically at 30°C for 3 days. Post incubation, the liquid fraction was removed by inverting the plates, and each well was washed three times with MilliQ water and air-dried. Subsequently, 100 µL of a 0.1% crystal violet solution (0.1% v/v crystal violet, 1% v/v methanol, and 1% v/v isopropanol in MilliQ water) was added to each well, followed by a 30-minute incubation at room temperature. After discarding the staining solution, wells were rinsed three times with MilliQ water. Biofilms were destained with 100 µL of a 30% acetic acid solution and incubated at room temperature for 30 minutes. OD_595_ of the destaining solution was measured for biofilm quantification.

#### Indole-3-Acetic Acid (IAA) assay

The protocol for IAA assay was adapted from Gilbert et al. (Gilbert et al., 2018). Briefly, the isolates were cultured in 5 ml R2A media, with and without 0.5 mg/mL L-tryptophan, for 3 days at 30°C. Following incubation, cultures were centrifuged at 14,000 rpm for 5 minutes, and 100 µl supernatant was transferred to a 96-well microplate. Subsequently, 200 µl of Salkowski reagent (2% of 0.5 M FeCl_3_ in a 35% HClO_4_ solution) was added to each well. IAA standards (0-10 µg/mL) were also prepared simultaneously. The microplate was incubated in the dark for 30 minutes, and absorbance values at OD_530_ was recorded to quantify IAA production using the standard curve.

#### ACC-deaminase activity

The method for measuring 1-Aminocyclopropane-1-carboxylate (ACC) deaminase production in bacterial isolates was adapted from the colorimetric ninhydrin assay described by Li et al. (Li et al., 2011). Briefly, SynCom isolates were cultivated in 2 ml of DF salinity minimum media supplemented with 3 mM ACC for 24 hours. Following incubation, 1 ml of the culture was centrifuged at 8,000 x g for 5 minutes. Subsequently, 100 μL of the resulting supernatant was diluted tenfold, and the ninhydrin assay, with ACC standards ranging from 0.5 mM to 0.005 mM, was used to quantify bacterial ACC consumption.

#### Phytate solubilization assay

Isolates were inoculated into 3 ml of filter-sterilized phytase-specific medium (PSM) (Hosseinkhani, Hosseinkhani, 2009) supplemented with 0.5 mM of Sodium phytate (Fisher Scientific). The cultures were then incubated for 48 hours at 30°C in a shaker incubator set at 80 rpm. Following incubation, the OD_600_ was measured, and the cultures were centrifuged at 10,000 rpm for 15 minutes. The resulting supernatant was retained for a colorimetric assay using the QuantiChrom™ Phosphate Assay Kit (BioAssay Systems, CA, USA). Absorbance was measured at 620 nm and compared against the provided phosphate standards included with the Phosphate Assay Kit.

#### Siderophore production assay

The capability of siderophore production by the isolates was assessed using the CAS overlaying medium assay (Acharya et al., 2023a; Louden et al., 2011; Pérez-Miranda et al., 2007). Briefly, each isolate was cultured on R2A agar plates at 30 °C for 3 days. Subsequently, blue CAS medium with 0.9% agar was overlaid onto the plates. The plates were then further incubated in the same conditions and monitored for color changes over an additional four days. The presence of a yellow halo surrounding colonies indicated positive siderophore production. As a positive control, *Burkholderia* sp. PA-E8, a known siderophore producer isolated at our laboratory, was included (Darling et al., 1998; Ong et al., 2016).

### Statistical analyses

#### Network analysis and SynCom member interactions

To understand the interspecies interactions from 0.1X R2A enrichments which have the highest diversity and most isolates, the top 50 most abundant ASVs among the core ASVs (present in >75% samples from the 4 generations) from this enrichment were selected. Their abundance matrix across all samples was analyzed using “NetCoMi” package (Peschel et al., 2021) in R. Pearson correlation coefficients greater than 0.3 and Student’s t-test results with p-values less than 0.05 were used to generate a sparse matrix for the network analysis. In the correlation network, each node represents an individual ASV, with different colors represent the corresponding modules. The edges connecting the nodes indicate a strong and significant correlation between the ASVs. The clustering in this network was used to group ASVs into modules that are densely interconnected internally but have sparse connections with other modules.

#### SynCom strain abundances in the rhizosphere under drought and salinity stress

To identify the changes in individual SynCom member’s abundances under stress, natural log fold differences between the control versus drought, rewatered drought, and salinity conditions were assessed using Analysis of Compositions of Microbiomes with Bias Correction (R package “ANCOMBC”) (Lin, Peddada, 2020; Mandal et al., 2015). False discovery rates were controlled using the Benjamini-Hochberg method.

#### Rhizosphere functional redundancy of SynCom members in stress conditions

To test whether the plants under stress modulate rhizosphere microbiome towards similar functions, functional redundancy (FR) across different environments (de Bello et al., 2021) estimated and compared using the R package “SYNCSA” (Debastiani, Pillar, 2012).

## Results

### Isolation of diverse rhizosphere bacteria from enrichments originating from Brachypodium rhizobiome

This study builds upon two prior studies where we first grew *Brachypodium* in natural soil to collect root microbiome (Acharya et al., 2023b) and subsequently enriched the community with various root exudate carbon substrates, resulting in 750 unique enrichments (Chen et al., 2024). In this study, we selected a few enrichments based on their high species richness, as indicated by Shannon diversity index, to isolate pure microbial strains. Selected enrichments included enrichments obtained on 0.1X R2A media, RCH2 media amended with Glutamine, and RCH2 media amended with mixed carbon. To obtain isolates, these enrichments were streaked on the corresponding agar media plates and incubated at 30°C in the dark for 7 days, yielding a total of 175 bacterial isolates. Taxonomic classification of the 16S rRNA genes for these isolates, based on SILVA (Quast et al., 2013), revealed that they belong to the following phyla: Alphaproteobacteria (15), Gammaproteobacteria (124), Actinobacteria (15), Firmicutes (13), Bacteroidetes (7) and Acidobacteria (1).

### Selection of isolates to construct a 15-member SynCom

To build a SynCom that is truly representative of the *Brachypodium* rhizosphere microbiome, we considered the following criteria: (1) relative abundance of different microbial taxa within the enrichments, (2) representation of phylogenetic diversity among the most abundant microbes, and (3) network interactions among the taxa. Based on these criteria, 15 isolates were selected from the cultured isolates to assemble the SynCom. These 15 selected isolates constituted a substantial 79.93% of the total ASVs in the 0.1X R2A enrichments and represented five bacterial phyla. Network analyses performed with the top 50 ASVs from this enrichment revealed six distinct interaction modules. In the final assembled SynCom, we included 6 strains from Module 1, 5 strains from Module 3, 3 strains from Module 4, and 2 strains each from Module 5 and Module 6 (Figure 1). The SynCom strains represent most of the network modules, and some members such as *Mesorhizobium* sp. RCC-202 (degree = 8) and *Leifsonia* sp. RCC-180 (degree = 5) have high network degrees (number of significant correlations connecting to the node), demonstrating their key role in maintaining the network structure. All but one i.e., *Bosea* sp. RCC-152.1, demonstrated a positive connection with at least one other strain across the different modules. Notably, the assembled SynCom also includes a strain of genus *Terriglobus*, known to be one of the key underexplored rhizosphere bacteria that play a crucial role in soil ecology (Eichorst et al., 2007, 2018; Kalam et al., 2022; Männistö et al., 2011; Pascual et al., 2015), yet are notoriously difficult to culture.

**Figure 1:**
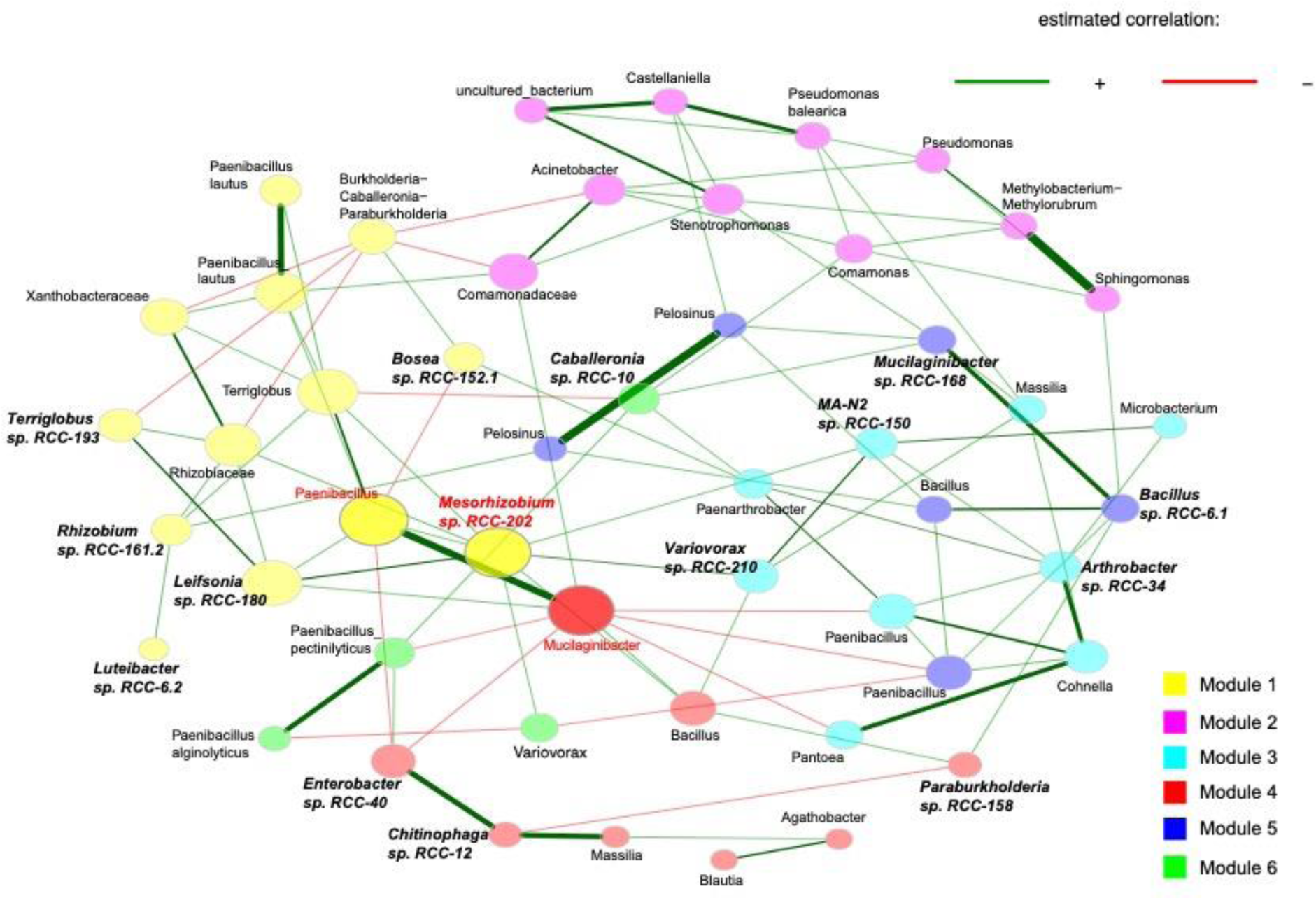
Association networks calculated using the 50 most abundant bacterial ASVs from 0.1X R2A enrichments. The abundance matrices from all samples were loaded into R to construct and visualize the network using the “NetCoMi” package. Networks were generated using the fast greedy clustering algorithm with a Pearson correlation coefficient threshold of ± 0.3, and a t-test (< 0.05) for sparse matrix generation. Eigenvector centrality is used for defining hubs and scaling node sizes. Positive correlations are displayed in green and negative in red. Shape color represents species clusters that are more likely to co-occur with one another than with species outside these modules. Hubs are highlighted in red with corresponding taxon names, whereas SynCom isolates have bolded and italicized names.

### Genomes of SynCom isolates encode multiple PGP traits

Genomic analysis of nearly complete genomes for all 15 isolates showed that they exhibited a wide range of genome sizes (3.7 Mbp -8 Mbp), GC content (35%-69%), and numbers of genes (3343-7740). The key general genome features of the SynCom isolates are outlined in Table S1. Genome-based taxonomic classification using the Genome Taxonomy Database (GTDB) (Parks et al., 2018), which employs 120 housekeeping marker genes for taxonomic classification, showed that the SynCom isolates belong to five phyla: Proteobacteria, Bacteroidota, Acidobacteriota, Actinobacteriota and Firmicutes (Figure 2).

**Figure 2:**
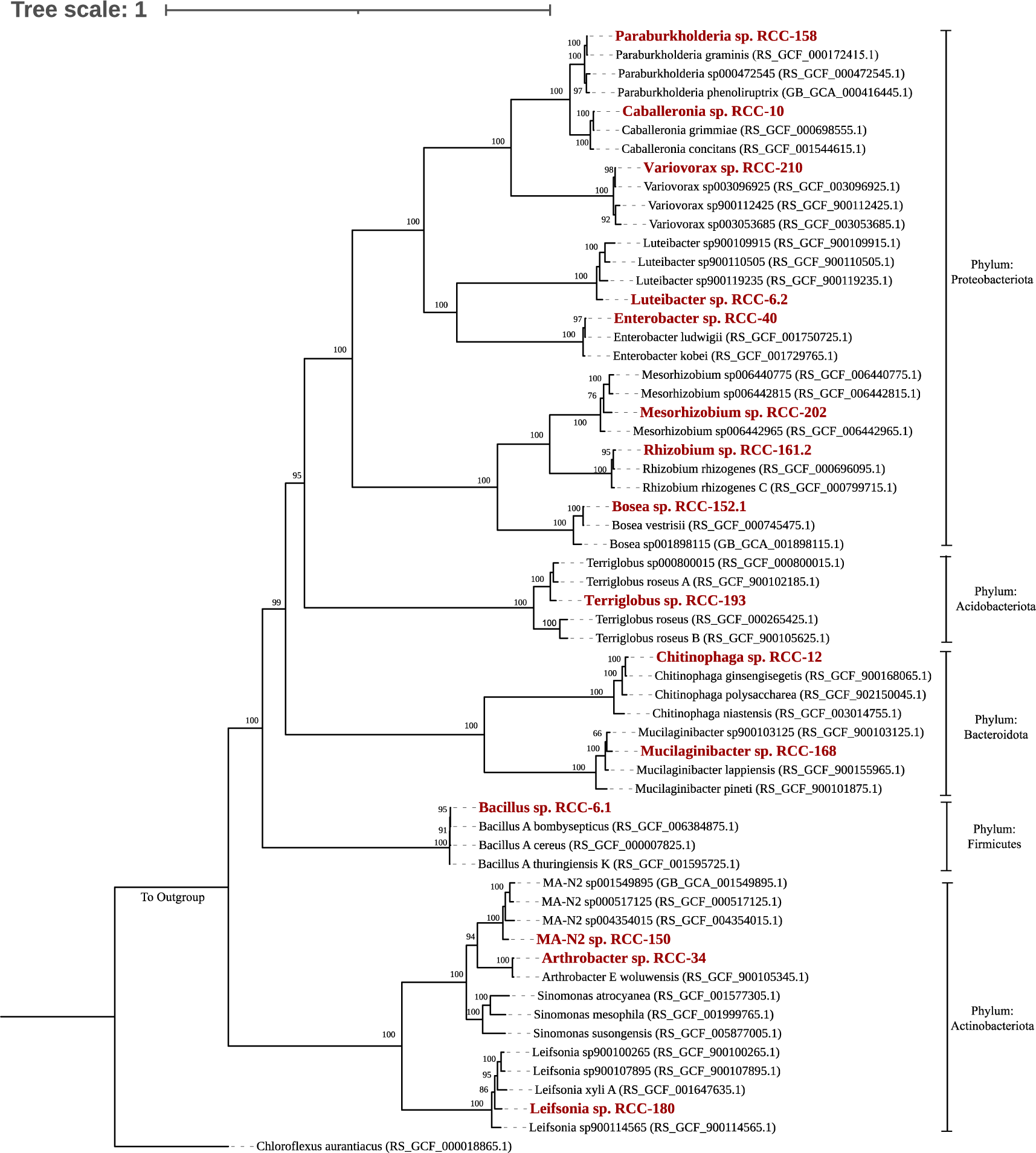
A phylogenomic tree for 15 SynCom isolates was constructed using multiple sequence alignment (MSA) from the concatenation of 120 marker proteins obtained via GTDB-tk workflow. The tree was visualized and edited using iTOL. To create the tree, 2-5 genomes closest to the SynCom isolates were included, with Genbank accession numbers in parentheses. The 15 SynCom members are highlighted in bold red. Phylum-level taxonomy for each genome is indicated on the right side of the tree. Bootstrap values (from 100 replicates) are displayed for nodes over 40 bootstrap supports. *Chloroflexus auranticus* was used as an outgroup to root the tree.

Most SynCom isolates encoded either a complete set of genes or the essential genes involved in pathways known to be associated with PGP traits that potentially contributes to overall plant growth (Table 1) (Bhattacharyya, Jha, 2012; Bruto et al., 2014; Gupta et al., 2014; Olanrewaju et al., 2021). Some of the important PGP traits encoded in the genomes of multiple SynCom isolates included 1-aminocyclopropane-1-carboxylate (ACC) deaminase production (9 isolates), Indole-3-acetic acid (IAA) for auxin synthesis (2 isolates), synthesis of antimicrobial compound phenazine (14 isolates) and gamma-aminobutyric acid (GABA) (12 isolates), genes associated with plant hormones like amidases (13 isolates), synthesis of volatile organic compounds (VOCs) i.e., 2,3 butanediol and acetoin (14 isolates), trehalose synthesis (13 isolates) and siderophore production or transportation (11 isolates) (Table 1). Among the 34 common PGP traits that we examined in this study, genomes of *Rhizobium* sp. RCC-161.2 and *Mesorhizobium* sp. RCC-202 stood out as they encoded the highest number of PGP traits, 16 and 14 respectively. The presence of these genes linked to various PGP mechanisms suggests that the SynCom strains could have beneficial effects on the plants.

**Table 1:**
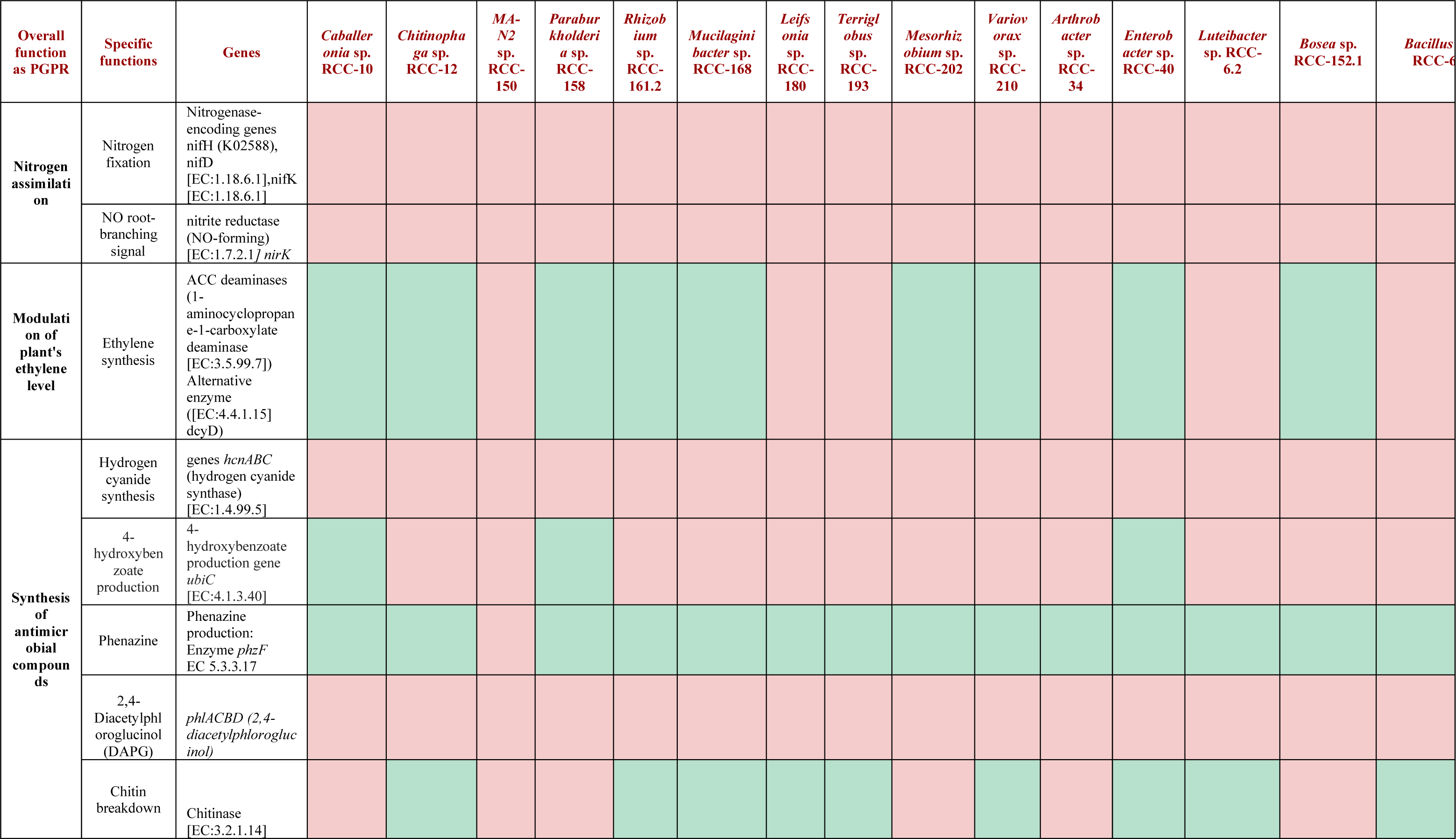
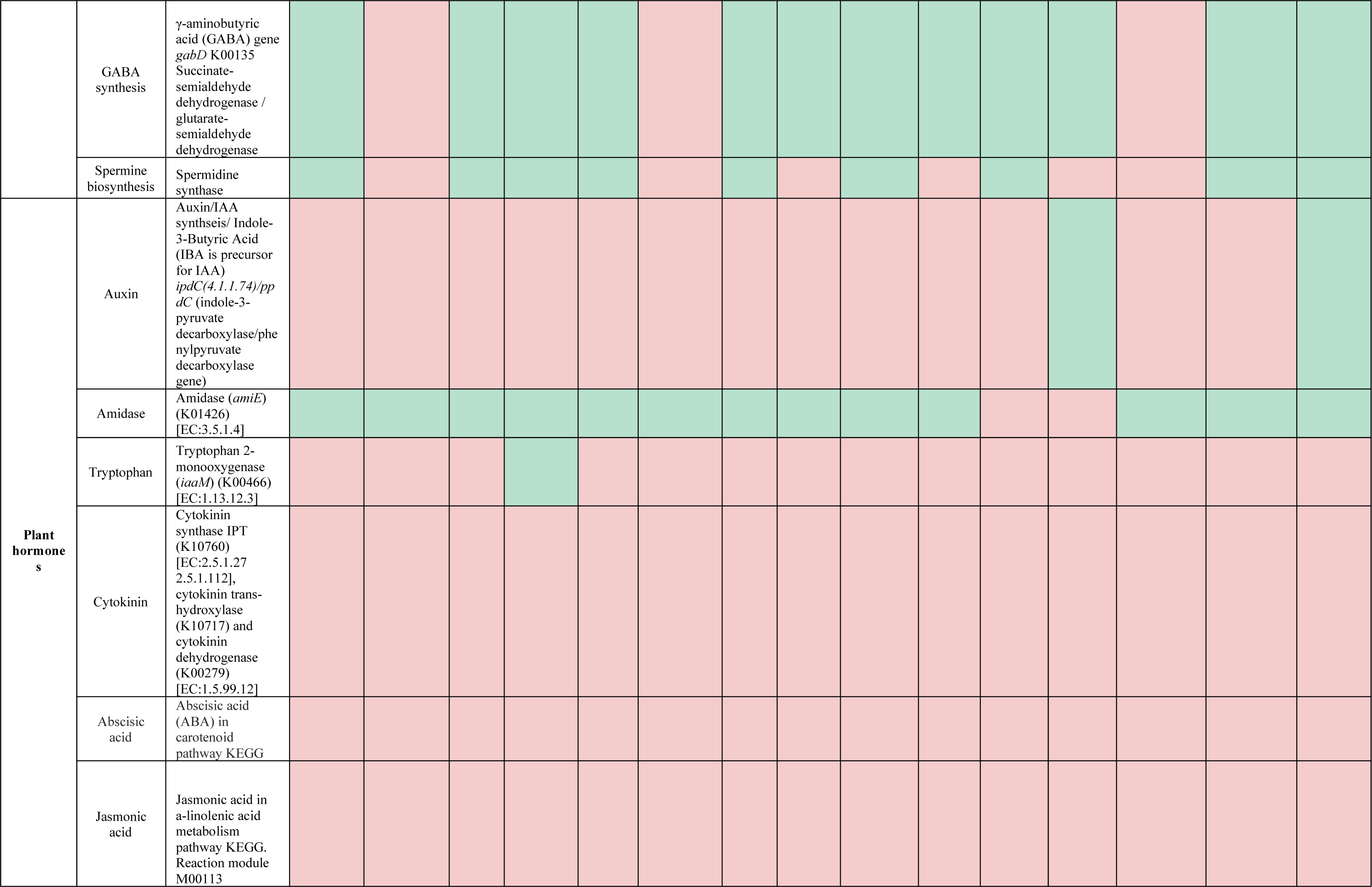
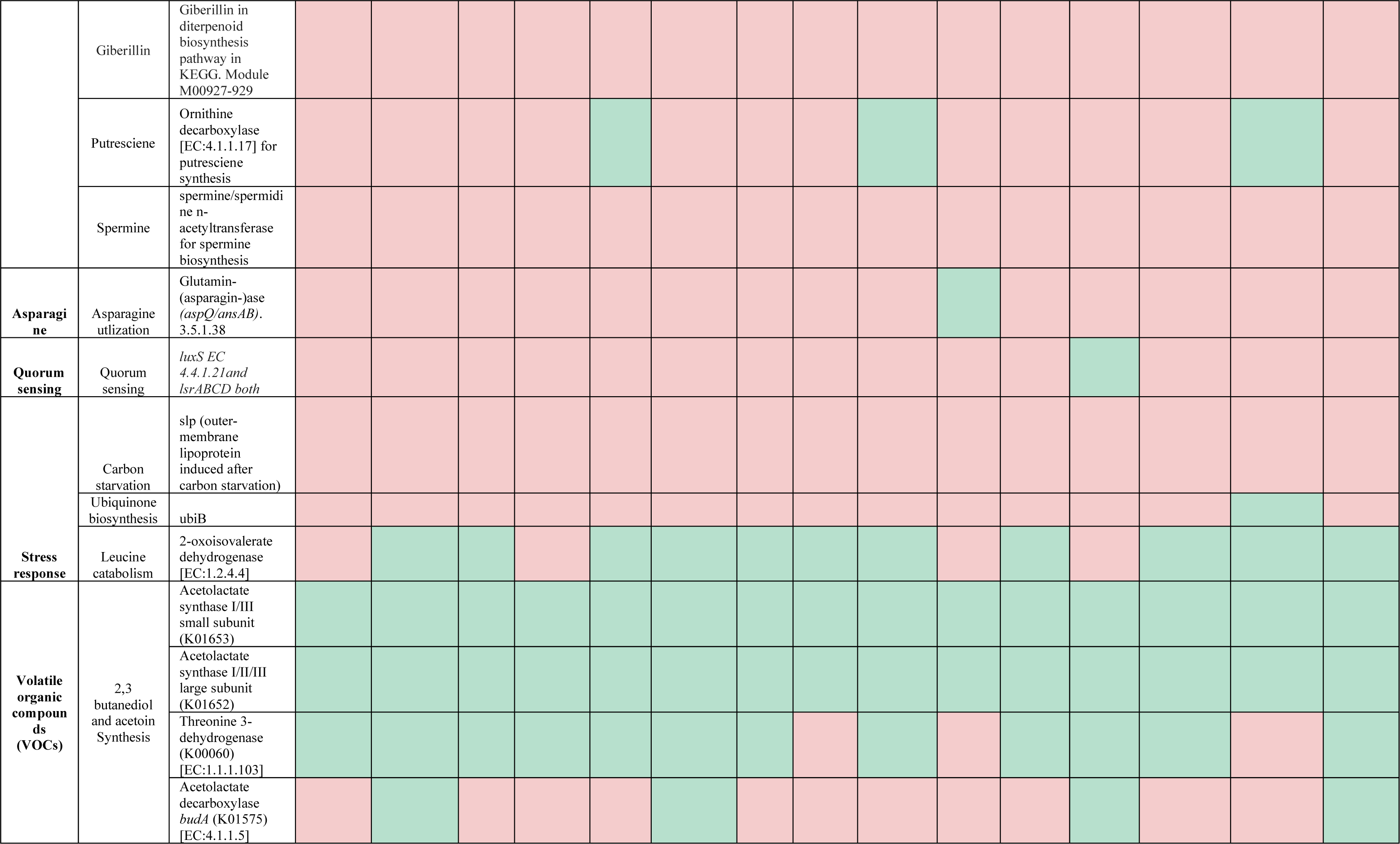
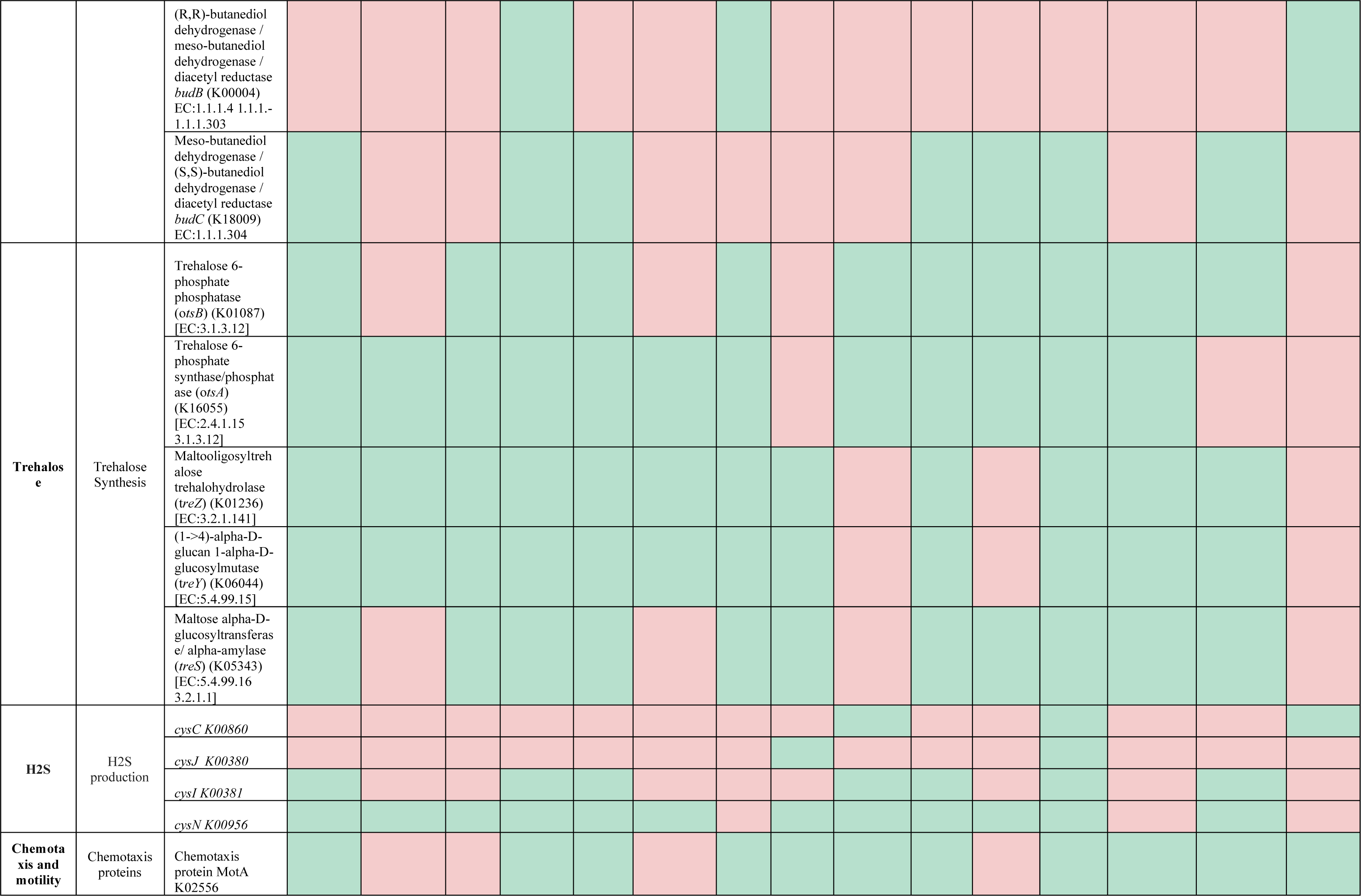
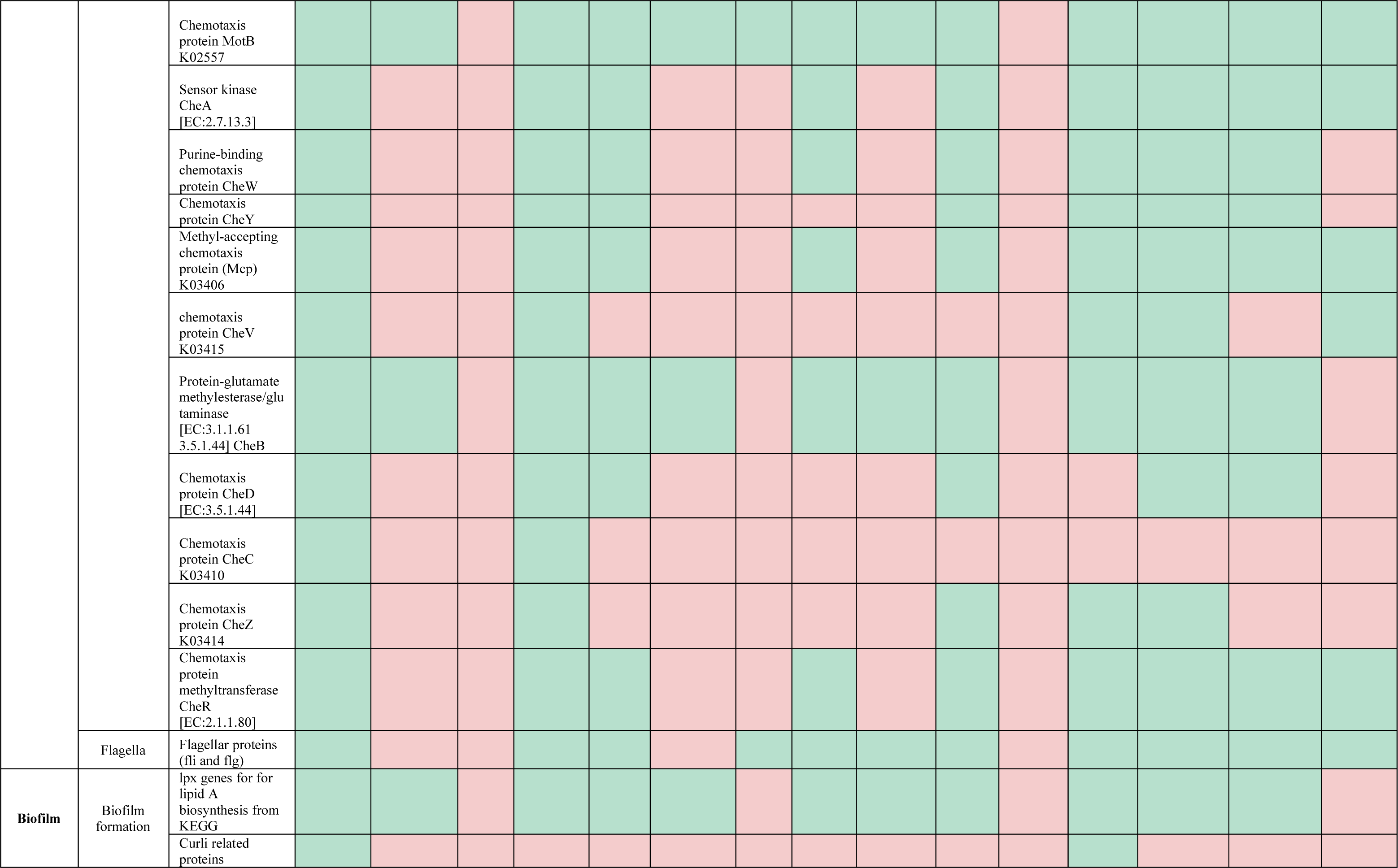
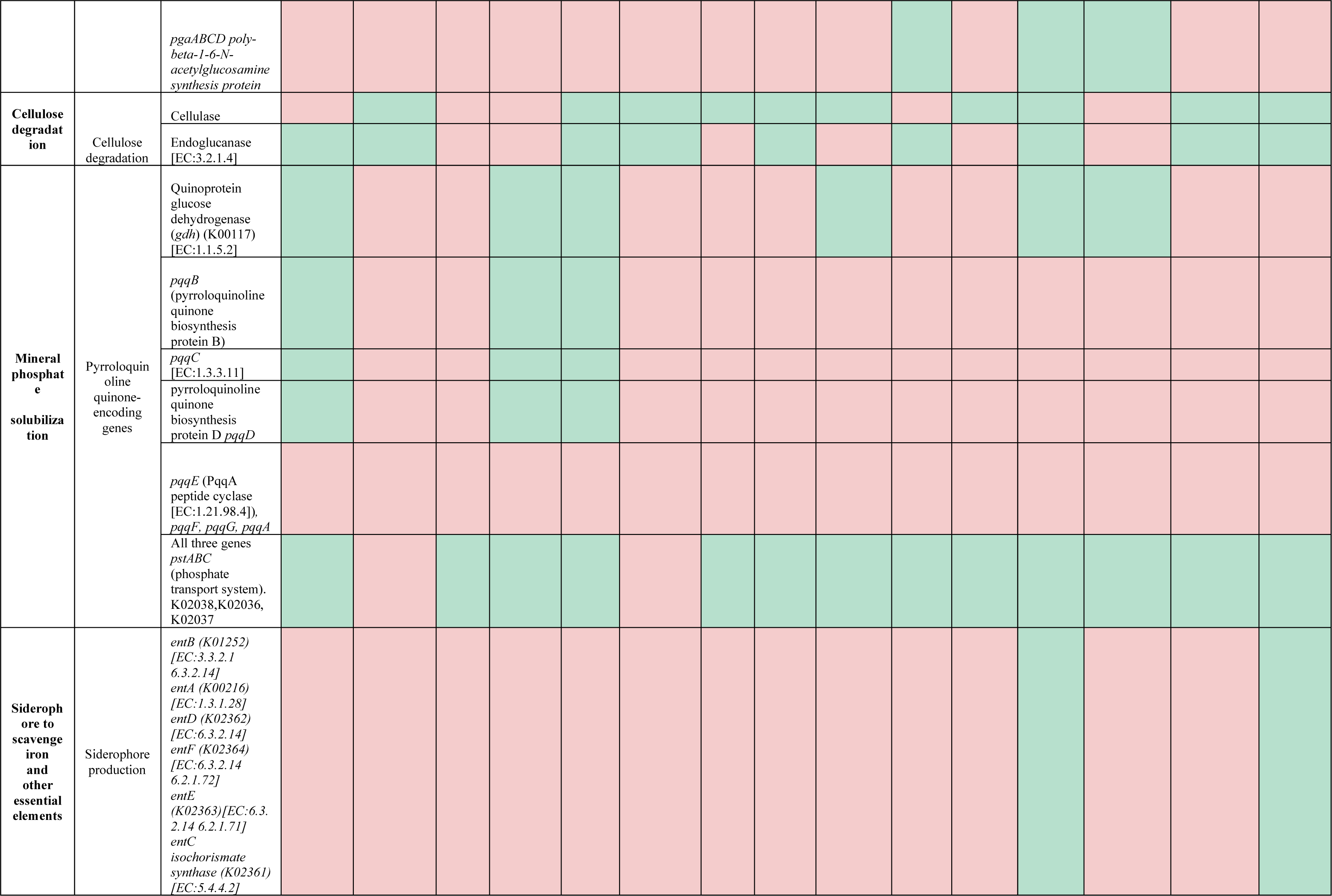
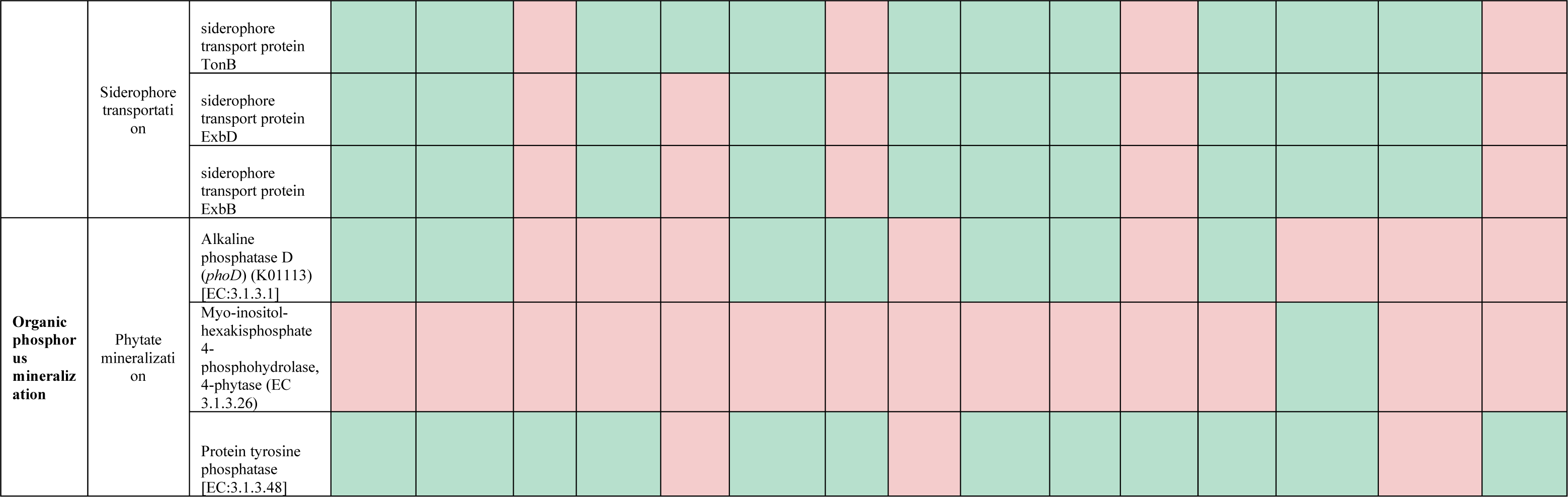
Plant Growth Promoting (PGP) factors encoded in genomes of SynCom isolates: The table categorizes genes according to their general Plant Growth Promoting Rhizobacteria (PGPR) functions (Column A), specific pathways (Column B), and the presence of these genes in the syncom member genomes (Column C). Genes found in the SynCom genomes are highlighted in green, while those that are absent are marked in red.

### SynCom strains express multiple PGP traits during phenotypic characterization

To validate the findings from genomic analysis for PGP traits in the isolates, we conducted phenotypic assays for five PGP traits: IAA production, ACC deaminase activity, phytate solubilization, siderophore production, and biofilm formation and results are summarized in Table 2. Six SynCom isolates demonstrated the ability to produce IAA when cultured with L-tryptophan with the produced IAA amount ranging from 0.25 to 23 µg/ml. Two strains, *Enterobacter* sp. RCC-40 and *Mucilaginibacter* sp. RCC-168, showed the highest IAA production levels of ∼23 µg/ml, higher when compared to the IAA production level of ∼5-7 µg/ml (Table 2) observed in multiple IAA-producing bacterial strains in the study by Gilbert et al. (Gilbert et al., 2018). All SynCom isolates exhibited ACC deaminase production, with nine strains capable of degrading more than half of the supplied 3 mM ACC, indicating a positive ACC deaminase activity. Five isolates including *Luteibacter* sp. RCC-6.2, *Terriglobus* sp. RCC-193, *Mucilaginibacter* sp. RCC-168, *Leifsonia* sp. RCC-180, and *Rhizobium* sp. RCC-161.2 demonstrated the ability to solubilize phytate, an organic form of phosphorus found in soils, with solubilization levels ranging from 0.0003 to 0.006 mg/dL (Table 2). Seven isolates could produce hydroxamate type siderophores acting as iron chelators, as confirmed by qualitative faint yellow-colored halo formation around the isolates colonies on a plate overlaid with chromeazurol S (o-CAS) medium (Pérez-Miranda et al., 2007) (Table 2). Lastly, crystal violet assay for testing biofilm formation in MS media demonstrated that nine SynCom strains exhibited a statistically significant biofilm-forming potential compared to the control (Table 2). Interestingly, *Mucilaginibacter* sp. RCC-168 tested positive for all five PGP traits tested here.

**Table 2:** Results summarizing the phenotypic assays for the plant growth-promoting (PGP) traits in SynCom isolates. A) ACC deaminase production: Values here indicate the quantity of ACC degraded (in mM) by the individual isolates when cultivated in media supplemented with 3 mM ACC. Darker green highlights indicate higher levels of ACC deaminase production. 2) Phytate Solubilization: The values presented here signify the amount of phytate consumed (in mg/dL) by each bacterium during their growth in phytate-specific media containing 0.5 mM Sodium phytate. Darker colors indicate greater capacities for phytate solubilization, while white color denotes a lack of phytate solubilization. 3) IAA production: Amount of IAA produced by the individual isolates when cultured in media containing 0.5mg/mL L-tryptophan and subjected to the Salkowski calorimetric test. Cells filled with darker colors signify higher capabilities for IAA production, while white color indicates a lack of IAA production. 4) Biofilm formation: Average OD595 values of the wash solution at the end of the biofilm assay of SynCom strains in MS media. The values for OD for the strains that had a significantly higher OD than the negative control (0.216) are listed here. Darker highlights correlated to higher biofilm formation. 5) Siderophore Production: (+) denotes positive siderophore production by the SynCom isolates characterized by the formation of a halo surrounding bacterial cultures upon overlaying with a blue-colored solution after 4 days of growth in a qualitative plate assay. Blank cells indicate the absence of siderophore production.

| SynCom Isolates | ACC consumed (mM) | Phytate Solubilization (mg/dL) | IAA production (ug/ml) | Biofilm formation (OD595) | Siderophore Production |
| --- | --- | --- | --- | --- | --- |
| *Bosea* sp. RCC-152.1 | 2.25 |  |  | 0.282 | + |
| *Luteibacter* sp. RCC-6.2 | 2.03 | 0.0003 |  | 0.337 |  |
| *Bacillus* sp. RCC-6.1 | 2.01 |  |  | 0.269 | + |
| *Enterobacter* sp. RCC-40 | 1.97 |  | 23.32 | 0.898 | + |
| *Paraburkholderia* sp. RCC-158 | 1.75 |  |  |  | + |
| *Caballeronia* sp. RCC-10 | 1.72 |  |  | 0.359 |  |
| *Arthrobacter* sp. RCC-34 | 1.64 |  | 8.30 | 0.736 |  |
| *Variovorax* sp. RCC-210 | 1.61 |  |  | 0.469 |  |
| *MA-N2* sp. RCC-150 | 1.57 |  |  |  |  |
| *Terriglobus* sp. RCC-193 | 1.56 | 0.0026 | 8.83 |  |  |
| *Chitinophaga* sp. RCC-12 | 1.40 |  |  |  | + |
| *Mesorhizobium* sp. RCC-202 | 1.39 |  | 0.25 | 0.259 |  |
| *Mucilaginibacter* sp. RCC-168 | 1.34 | 0.0043 | 23.32 | 0.316 | + |
| *Leifsonia* sp. RCC-180 | 1.31 | 0.0065 |  |  |  |
| *Rhizobium* sp. RCC-161.2 | 1.17 | 0.0026 | 3.13 |  | + |

### SynCom strains exhibit persistence in vitro and in planta

Ensuring the SynCom’s stability and its effective colonization on the plants grown in conditions mimicking natural habitat is a crucial element in SynCom development and deployment (Liu et al., 2022; Pradhan et al., 2022; Shayanthan et al., 2022). We assessed the stability of SynCom both in the presence and the absence of plants. In the *in vitro* experiment, where equal cell number of SynCom strains were grown as a consortium and subcultured in 0.2X MS media for 3 weeks, the 16S rRNA gene sequence analysis showed that 14 of the 15 strains continued to persist by the end of the third week. However, four strains (*Arthrobacter* sp. RCC-34*, Leifsonia* sp. RCC-180*, Luteibacter* sp. RCC-6.2, *and Mesorhizobium* sp. RCC-202) exhibited very low relative abundances of <0.1% and *Bacillus* sp. RCC-6.1 was seemingly absent by the third week (Figure 3A). Conversely, members such as *MA-N2* sp. RCC-150, *Paraburkholderia* sp. RCC-158 and *Chitinophaga* sp. RCC-12, dominated the culture with an average relative abundance of 60%, 26% and 8%, respectively (Figure 3A, Supplementary Figure S1). To further probe into SynCom’s stability, we conducted a leave-one-out experiment, where the overall growth in 0.1X R2A broth media was analyzed for multiple 14-membered SynCom created by excluding one SynCom member at a time (Figure 3B). The results indicated that all these 14-membered consortia grew as well or better than the 15-membered SynCom, demonstrating the stability of these consortia and the lack of any negative interactions between SynCom strains (Figure 3B).

**Figure 3:**
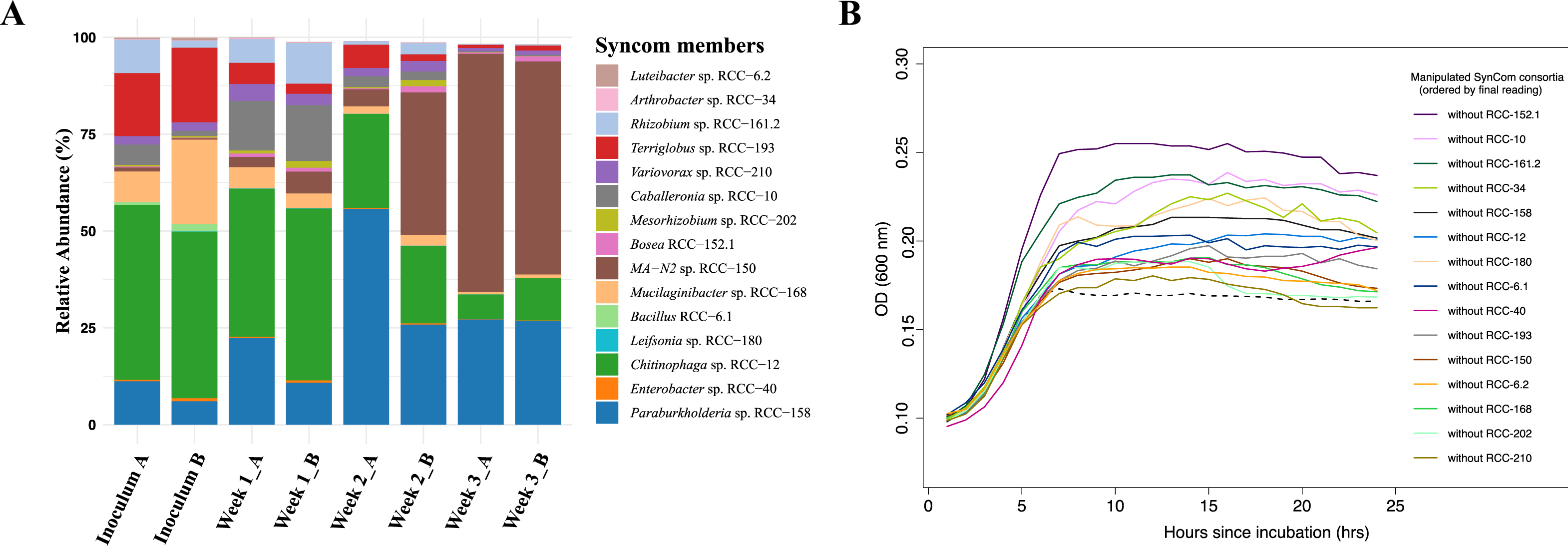
Stability of Syncom members: 3A) Changes in the microbial community over three weeks when the 15-member SynCom is cultured in 0.2X MS media and transferred weekly. The initial bar on the x-axis reflect the relative abundance of 15 SynCom members in the Inoculum, followed by pairs of bars representing duplicates for samples taken at the end of Weeks 1, 2, and 3. Y-axis shows the relative abundances of each SynCom strain which is represented by different color as shown in the legend. 3B) Growth dynamics measured by average OD_600_ over 24 hours, for various manipulated SynCom mixes from the leave-one out experiment in 0.1X R2A broth (n = 3). The dotted curve represents the bacterial mix with all 15 SynCom members. Other colored curves represent bacterial mixes with one SynCom member omitted, with the omitted members indicated in the legend.

To assess *in planta* stability of the SynCom, we added the inoculum containing equal cell numbers (∼ 4x10^7^ cells) of each SynCom strains, to the base of the *Brachypodium* seedlings in the pots and grew them for 3 weeks (Supplementary Figure S2). The relative abundance of the strains at the end of 3 weeks showed that multiple strains were highly enriched with relative abundance of more than 1%. These include *Paraburkholderia* sp. RCC-158 (34%), *Enterobacter* sp. RCC-40 (15%), *Chitinophaga* sp. RCC-12 (13%), *MA-N2* sp. RCC-150 (7%), and *Luteibacter* sp. RCC-6.2 (3%) (Supplementary Figure S1). Notably, in both the *in vitro* and the *in planta* experiments, three strains i.e., *Paraburkholderia* sp. RCC-158, C*hitinophaga* sp. RCC-12, and *MA-N2* sp. RCC-150 emerged as the dominant strains, comprising more than half of the total microbial community (Figure 3A, Figure 4B). Besides these, several other SynCom strains achieved higher relative abundances in the *in planta* experiment compared to their relative abundances in the *in vitro* experiment. This enrichment of multiple SynCom strains within the plant rhizobiome is likely driven by the diverse carbon sources exuded from plant roots, enabling beneficial microbes to thrive (Vives-Peris et al., 2020; Wen et al., 2022). The detectable presence of all SynCom strains within the rhizobiome after 4 weeks of plants growth provides further stability of this SynCom when grown with the plant.

**Figure 4:**
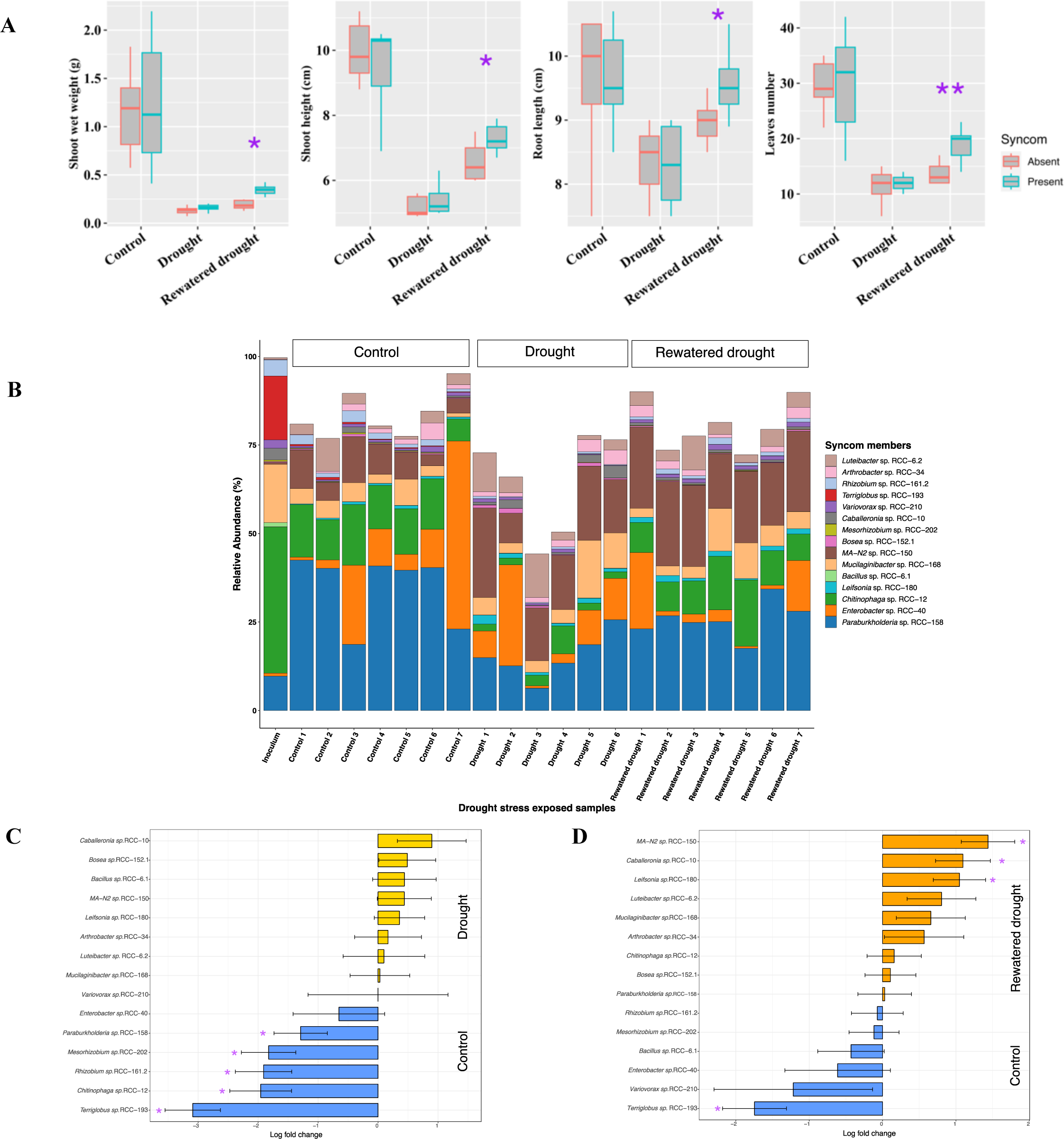
Figures depicting the plant phenotypic traits and microbial 16S r RNA gene abundances profile from the SynComdrought stress experiment. The experiment comprised three sets: the Control set, Drought set, and Rewatered Drought set. A) Plant phenotype results: Comparisons for the shoot weight, shoot height, root length, and leaves numbers in *Brachypodium*, with and without SynCom members, during a 3-week exposure to drought stress. Each experimental category included seven *Brachypodium* plants. Significant values determined by t-tests are indicated by purple asterisks positioned above the corresponding bars. B) 16S rRNA gene abundances profiles: In the barplot, each bar represents a replicate (shown in x-axis) in the corresponding experimental set as indicated above the bars. Y-axis shows the relative abundances values for e ach SynC om isolates represented by different color in legend, while all other bacterial groups are combined into the “Others” group, depicted inwhite on the plot. It is important to note that certain DNA samples in the experimental sets i.e., rewatered drought encountered sequencing issues, leading to a reduced sample size of 5. 4C) The natural log fold changes (lfc) for SynCom strains in stress conditions, as measured by “ANCOMBC” wtih drought (yellow) and 4D) rewatered drought (orange), when compared to the SynCom-am ended control plants (blue). On y-axis, negative values indicate that taxa are more abundant in control conditions, while positive values show greater abundance in drought or rewatered drought conditions. Bars indicate changes in the natural log fold, and error bars represent standard errors. Asterisks indicate significant changes inthe natural log fold after the Benjamini-Hochberg adjustment of the false discovery rate (p < 0.05).

### Drought and salinity stress in Brachypodium enrich specific SynCom strains

We evaluated the efficacy of the SynCom on the plant phenotype, after subjecting the plants to common environmental stresses i.e., drought and osmotic stress for three weeks. To simulate drought stress, plants received watering equivalent to 40% of their saturation weight. These plants exhibited no significant differences in shoot height, shoot weight, root length, and leaf number between SynCom-amended and unamended groups (Figure 4A). Root microbiome analysis, based on 16S rRNA gene sequences, revealed that all SynCom strains persisted (Figure 4B). However, a significant (p-value <0.05) decrease in log fold change (lfc) in relative abundances of *Paraburkholderia* sp. RCC-158 (lfc=1.29), *Rhizobium* sp. RCC-161.2 (lfc=1.91), *Mesorhizobium* sp. RCC-202 (lfc=1.82), *Terriglobus* sp. RCC-193 (lfc=3.09) and *Chitinophaga* sp. RCC-12 (lfc=1.96) was observed in SynCom-amended plants under drought conditions compared to control sets (Figure 4C).

To mimic drought and rewetting conditions in nature, a ‘rewatered drought’ condition was implemented, where drought-treated plants were subsequently watered similar to the control condition for one week before sampling. Under rewatered drought conditions, SynCom-amended plants demonstrated a significant improvement in all measured plant phenotype parameters i.e., shoot weight, shoot height, root length and leaves number, when compared to the unamended plants (Figure 4A). Root microbiome analysis revealed that all SynCom strains persisted. There was a significant decrease in *Terriglobus* sp. RCC-193 (lfc = 1.74), and a significant increase in *MA-N2* sp. RCC-150 (lfc=1.4), *Leifsonia* sp. RCC-180 (lfc=1.04) and *Caballeronia* sp. RCC-10 (lfc=1.09), when compared to control sets (Figure 4D).

Lastly, to simulate salinity stress, plants received the watering equivalent supplemented with 60 mM sodium chloride (NaCl). SynCom-amended plants during this stress exhibited a significant improvement in shoot height when compared to unamended plants (Figure 5A). Similar to the drought stress results, all SynCom strains persisted under salinity stress (Figure 5B). However, no significant changes in the relative abundance of any SynCom strains were observed (Figure 5C).

**Figure 5.**
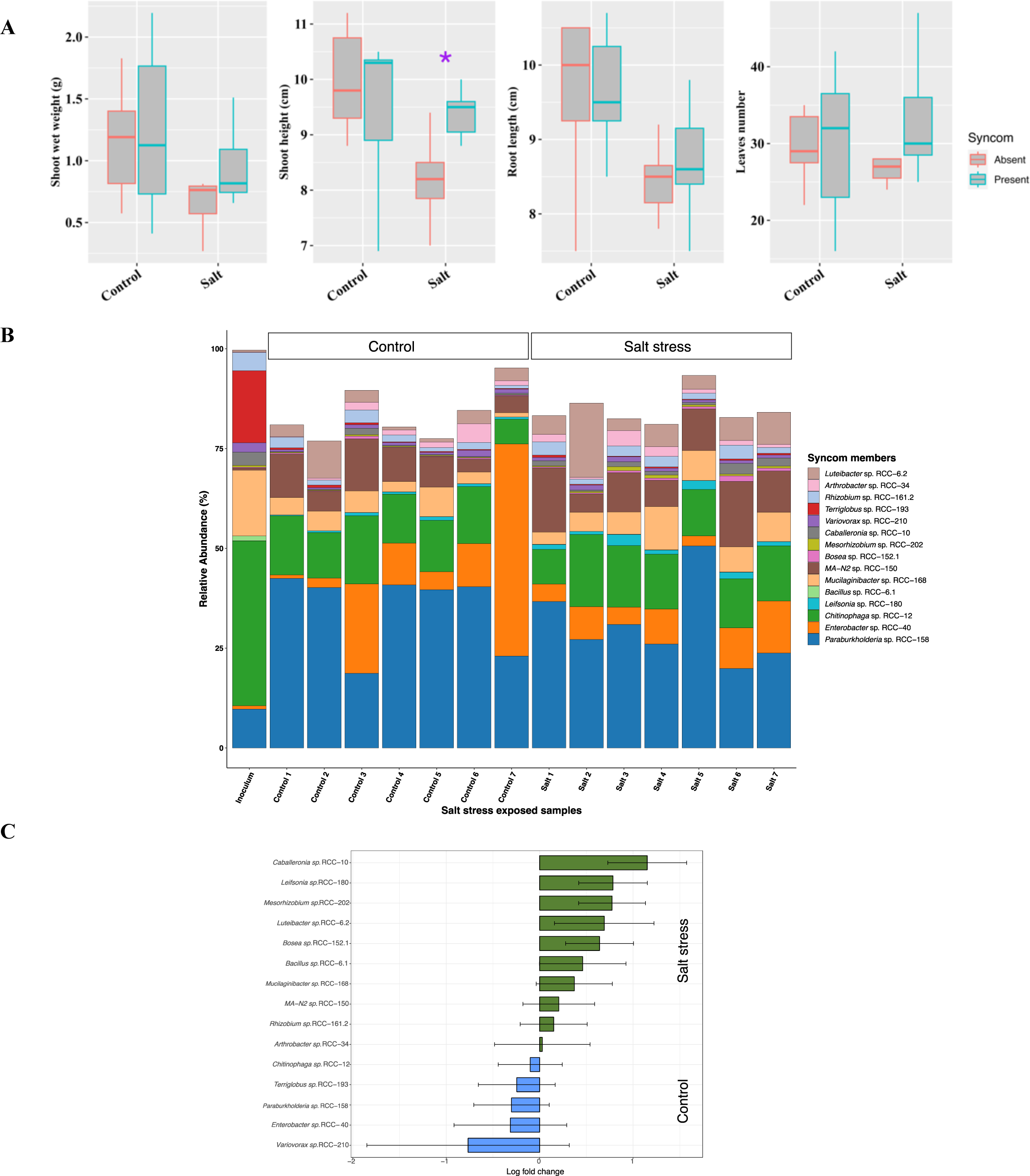
Figures depicting the plant phenotypic traits microbial 16S rRNA gene abundances profile from the SynCom salt stress experiment. The experiment comprised two sets i.e, the Control set and the Salt stress. 5A) Plant phenotype result: Comparisons for the shoot weight, shoot height, root length and leaves numbers in *Brachypodium*, with and without SynCom members, during a 3-week exposure to salt stress. Each experimental category included seven *Brachypodium* plants. Significant values determined by t-tests are indicated by purple asterisks positioned above the corresponding bars. 5B) 16S rRNA gene abundance profiles: In the barplot, each bar represents a replicate (shown in x -axis) in the corresponding experimental set as indicated above the bars. Y-axis shows the relative abundances values for e ach Sync om isolate represented by different color in the legend, while all other bacterial groups are combined into the “Others” group, depicted in white on the plot. 5C) The natural log fold changes (lf c) for SynCom strains in stress conditions, as measured by “ANCOMBC” with salt stress (green) when compared to the SynCom-am ended control plants (blue). On y-axis, negative values indicate that taxa are more abundant in control conditions, while positive values show greater abundance in salt conditions. Bars indicate changes in the natural log fold, and err or bars represent standard errors. None of the SynC om members was found to have significant changes in natural log fold after the Benjamini -Hochberg adjustment of false discovery rate (p < 0.05).

### SynCom strains in the rhizosphere exhibit distinct spatial localization

Previous studies have highlighted that plants secrete specific compounds as root exudates to recruit select microbes, especially under stress conditions (Vives-Peris et al., 2020). Given that the exudates might vary between root tips and base (Dragišić Maksimović et al., 2021), we investigated the recruitment and spatial distribution of SynCom strains on the root. After 21 days of drought and salinity conditions, we sampled both the root tip and base separately for 16S rRNA amplicon sequencing to investigate colonization. Distinct spatial colonization was observed for specific SynCom strains. *MA-N2* sp. RCC-150 showed preferential colonization to the root tips under all tested conditions i.e., drought, rewatered drought and salinity stress, whereas *Leifsonia* sp. RCC-180 showed preferential colonization to the root tips only under rewatered drought conditions (Figure 6). Four strains i.e., *MA-N2* sp. RCC-150, *Mucilaginibacter* sp. RCC-168, *Variovorax* sp. RCC-210, and *Arthrobacter* sp. RCC-34 showed preferential colonization on the root base compared to the inoculum under control conditions (Figure 6).

**Figure 6:**
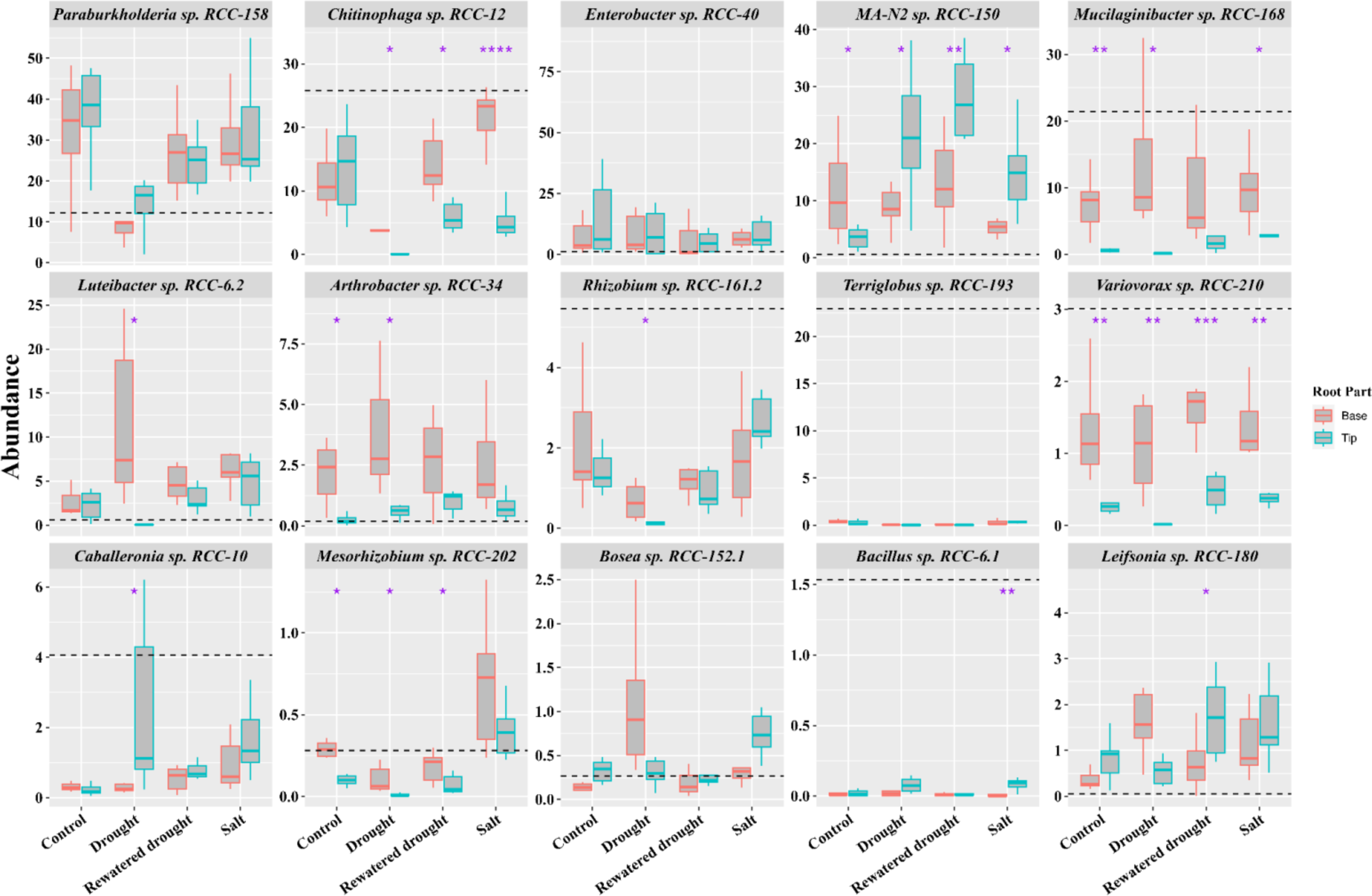
Spatial localization of SynCom strains in the *Brachypodium* root system: Boxplots show the 16S rRNA gene abundances for the SynCom isolates across the root tip and base in SynCom-amended plants under control and drought and salt stress conditions (as indicated on the x-axis) after 21 days. The y-axis displays the relative percentage abundances of the SynCom members indicated atop each boxplot panel. Significant differences determined by t-tests between the root parts (tip and base) within each condition are marked with an asterisk above and between the respective bars. The black horizontal dashed line indicates the percentage of strain abundance in the inoculum.

## Discussion

In this study, we developed a framework that integrated both culture-dependent and culture-independent methods, combining the strengths of reductionist and holistic approaches to create a SynCom. This combinatorial approach enabled us to assemble a well-defined SynCom comprising 15 bacterial genera from the *Brachypodium distachyon* rhizobiome. The 15-member SynCom stands out for several reasons. First, we used network analysis to examine the microbial interactions (both positive and negative) from the enrichments of the *Brachypodium* rhizobiome (Chen et al., 2024) (Figure 1), and assembled the SynCom guided by these interactions. This approach ensures that the SynCom is ecologically relevant and functionally intact (Toju et al., 2020). Notably, a keystone species identified through network analysis *Mesorhizobium* sp. RCC-202, is included in the SynCom which adds to the robustness of the community since keystone species are often responsible for shaping the microbial community structure and dynamics (Agler et al., 2016). Second, unlike most SynComs that focus on a few phyla like Proteobacteria, Actinobacteria, and Firmicutes (Marín et al., 2021), this SynCom also includes phyla Bacteroidetes and Acidobacteria (Figure 2), both of which are particularly relevant in plant rhizosphere (Kalam et al., 2022; van Overbeek, 2013), as the diverse species composition has been shown to increase genetic diversity, functional overlap, stress tolerance, and improve growth and productivity in the host plants (Coker et al., 2022; Flores-Núñez et al., 2023; Li et al., 2021; Timm et al., 2016).

Another key highlight of this work is the rigorous assessment of the stability and robustness of the SynCom under both *in vitro* and *in planta* conditions. Stability is a highly desirable trait for creating successful microbial consortium (McClure et al., 2020; Zegeye et al., 2019) for agricultural applications (Shayanthan et al., 2022). The 15-membered SynCom showed remarkable stability both in the *in vitro* experiment where 14 out of the 15 members were detected (Figure 3A), and in the *in planta* experiments where all members were detected by the end of three weeks (Figures 4B, 5B). Notably, more strains were enriched with higher relative abundances in the *in planta* experiments when compared to the *in vitro* conditions (Supplementary Figure S1). This enrichment of diverse strains in the *in planta* experiments may be attributed to metabolite and nutrient-rich root exudates known to specifically modulate microbial abundance in the rhizosphere (Huang et al., 2019; Vives-Peris et al., 2020). An interesting observation was made in the *in vitro* experiments, where three out of the four Gram-positive bacterial genera such as *MA-N2* sp. RCC-150, *Leifsonia* sp. RCC-180, and *Arthrobacter* sp. RCC-34 demonstrated notable persistence over 3 weekly transfers of the SynCom in the MS media. *MA-N2* sp. RCC-150, in particular, showed an exceptionally high relative abundance (55%) at the end of this *in vitro* experiment (Supplementary Figure S2). This finding is noteworthy, as previous studies have reported that Gram-positive bacteria typically exhibit less persistence across multiple passages *in vitro* compared to their Gram-negative counterpart (Fonseca-García et al., 2024). This finding of persistent Gram-positive bacteria in the SynCom over multiple passages *in vitro* is important, as it might indicate similar persistence in the environment (Qi et al., 2022; Schimel et al., 2007), playing a significant role in aiding the host plant’s survival during stress conditions.

Additionally, while systematically examining the PGP traits of the SynCom members, we discovered discrepancies between genomic predictions and phenotypic expression of some tested PGP traits. For instance, laboratory experiments revealed higher frequencies of IAA production, ACC deaminase activity, phytate solubilization, siderophore production, and biofilm formation among the isolates than what genomic analysis had predicted (Tables 1, Table 2). This discrepancy may be attributed to challenges including insufficient gene annotation in existing literature, and the presence of alternative undefined pathways in some genomes (Agarwal, Shendure, 2020; Dimonaco et al., 2022; Gingeras, 2007; Kang et al., 2022; M A Basher et al., 2020; Sentausa, Fournier, 2013; Warren et al., 2010). Therefore, our findings also highlight the importance of integrating both genomic predictions and lab-based phenotype assays for validating PGP traits in bacteria.

We further evaluated whether SynCom could enhance *Brachypodium* growth under stress conditions such as drought and salinity, two abiotic stresses that limit global food crop production (Uddin et al., 2016). Under drought, rewatered drought, and salinity stress, SynCom-amended plants exhibited improved plant phenotype compared to the control plants without SynCom. However, statistically significant enhancement in plant phenotypes was only observed in SynCom-amended plants under rewatered drought conditions (Figures 4A, 5A). Notably, plants often do not exhibit immediate phenotypic changes despite microbial amendment (do Amaral et al., 2016), similar to our observed results. Additional transcriptome-based analyses are necessary to better understand the positive effects of SynCom members on plant growth, particularly during stress events.

Several changes related to microbial recruitment and their interaction under different environmental stress conditions were observed. In rewatered drought conditions, we found enhanced positive network interactions among SynCom members when compared to the control (Supplementary Figure S4), potentially promoting microbial interaction and function and benefiting the host plant (Layeghifard et al., 2017; Martins et al., 2023). This was reflected in higher plant shoot mass and root length (Figure 4A). The isolates that consistently showed higher relative abundance under rewatered drought conditions belong to genera known to alleviate drought stress in plants (Lim, Kim, 2013; Nordstedt et al., 2021; Platamone et al., 2023; Tallapragada et al., 2016), and many of them encoded genes for multiple osmoprotectant compounds (Supplementary Table S2). All strains except *Bacillus* RCC-6.1 encoded genes for trehalose biosynthesis (Table 1), a crucial mechanism enabling microbes to aid plants in managing drought and salinity stresses (Hassan et al., 2023; Kosar et al., 2018; Sharma et al., 2020). Additionally, many strains possess multiple sodium (Na^+^) and potassium (K^+^) transporters (Supplementary Table S3), which together with the osmoprotectants, help microbes maintain cellular homeostasis to resist osmotic stresses from salinity and drought conditions (Ahmad et al., 2022; Gunde-Cimerman et al., 2018). This capability potentially aids plants in coping with these abiotic stresses (Hao et al., 2021; Sharma et al., 2019; Singh et al., 2022). Lastly, all SynCom isolates showed some level of ACC deaminase activity (Table 2), which has been correlated to plant tolerance to salt stress (Orozco-Mosqueda et al., 2020). Interestingly, despite there being no significant enhancement in plant phenotype under drought stress (Figure 3A), we observed significantly higher bacterial functional redundancy signifying that the SynCom members enriched during drought stress encode more similar PGP traits (Table 1) as compared to the SynCom members enriched in control conditions (Supplementary Figure S3). The inconsistent results between drought and rewatered drought treatments indicate a plant growth boost associated with SynCom at the initial rewetting stage after the drought rather than during the drought. The higher functional redundancy under drought (Supplementary Figure S3) also suggested that plants may modulate rhizosphere bacterial community structure based on similar PGP functions rather than recruiting a few specific taxa (Louca et al., 2018; Ngumbi, Kloepper, 2016; Zia et al., 2021). This also suggests that the benefits of SynCom under drought may be more closely linked to enhancing plant survival rather than promoting growth (Louca et al., 2018; Ngumbi, Kloepper, 2016; Zia et al., 2021).

Microbial colonization at root tips and bases serves specific functions (Kamilova et al., 2005), often driven by the spatial gradients of root exudates. These patterns emphasize the intricate and mutually beneficial interactions between plants and their associated microbial communities (Birt et al., 2022; Wang et al., 2017; Wheatley, Poole, 2018). In our *in planta* experiments, we observed marked preference for colonization at the root tip under stress conditions (Figure 6) for Gram-positive bacteria *MA-N2* sp. RCC-150, *Leifsonia* sp. RCC-180, and *Bacillus* sp. RCC-6.1. This increase at the root tip, despite the initial inoculum being applied to the root base, suggests active migration of these strains which is supported by the presence of motility genes such as pili, flagella, or chemotaxis genes in their genomes (Table 1). Overall, the mutually beneficial interactions between plant and microbes could explain the localization patterns. On one hand, plants under stress may release specific compounds such as osmoprotectants, stress hormones, antioxidants, and certain metabolites (Mavrodi et al., 2021; Yoon et al., 2019). For instance, *Brachypodium* differentially regulates primary metabolites such as proline and soluble sugars, which are higher in drought-tolerant varieties (Shi et al., 2015). Given that the root tip is a center for active exudation and signaling molecules (Aufrecht et al., 2022), it is reasonable to hypothesize that these bacteria preferentially utilize stress-induced metabolites, influencing their localization pattern. On the other hand, numerous Gram-positive bacteria, including Actinobacteria and Bacilli, produce metabolites crucial for growth, stress tolerance, and plant defense mechanisms (Ek-Ramos et al., 2019; Grover et al., 2016). The colonization of these beneficial microbes near the root tip, a zone of active growth and nutrient absorption, could significantly contribute to plant sustenance under stress conditions. Besides, stress-induced alterations in root architecture, such as the development of deeper or more compact root systems, can modify the physical environment of the rhizosphere. These architectural changes indirectly affect water and nutrient distribution within microbial microhabitats (Jobbágy, Jackson, 2001; Nunes et al., 2020) thereby influencing microbiome assembly (Hartman, Tringe, 2019).

Lastly, while plant experiments integrated with genomic information of the SynCom are valuable for understanding plant phenotype and microbial abundances, incorporating transcriptomic and metabolomic data will be crucial to delve deeper into the underpinnings of plant-microbial interactions when exposed to stress as well as upscaling the utilization of SynCom in agricultural setting.

## Acknowledgements

This material by m-CAFEs Microbial Community Analysis & Functional Evaluation in Soils, a Science Focus Area led by Lawrence Berkeley National Laboratory is based upon work supported by the U.S. Department of Energy, Office of Science, Office of Biological Environmental Research under contract number DE-AC02-05CH11231.

## Data availability

The 16S rRNA amplicon sequences and assembled genomes for the SynCom isolates as well as the raw datasets for the stress experiments are available in the NCBI repository, under the BioProject PRJNA1127609. The KBase narrative containing the genome assemblies and annotation tools run on the assemblies are also publicly available at <u>narrative.kbase.us/narrative/135275</u>.

