## Supplementary figures and tables for "A stable 15-member bacterial SynCom promotes *Brachypodium* growth under drought stress"

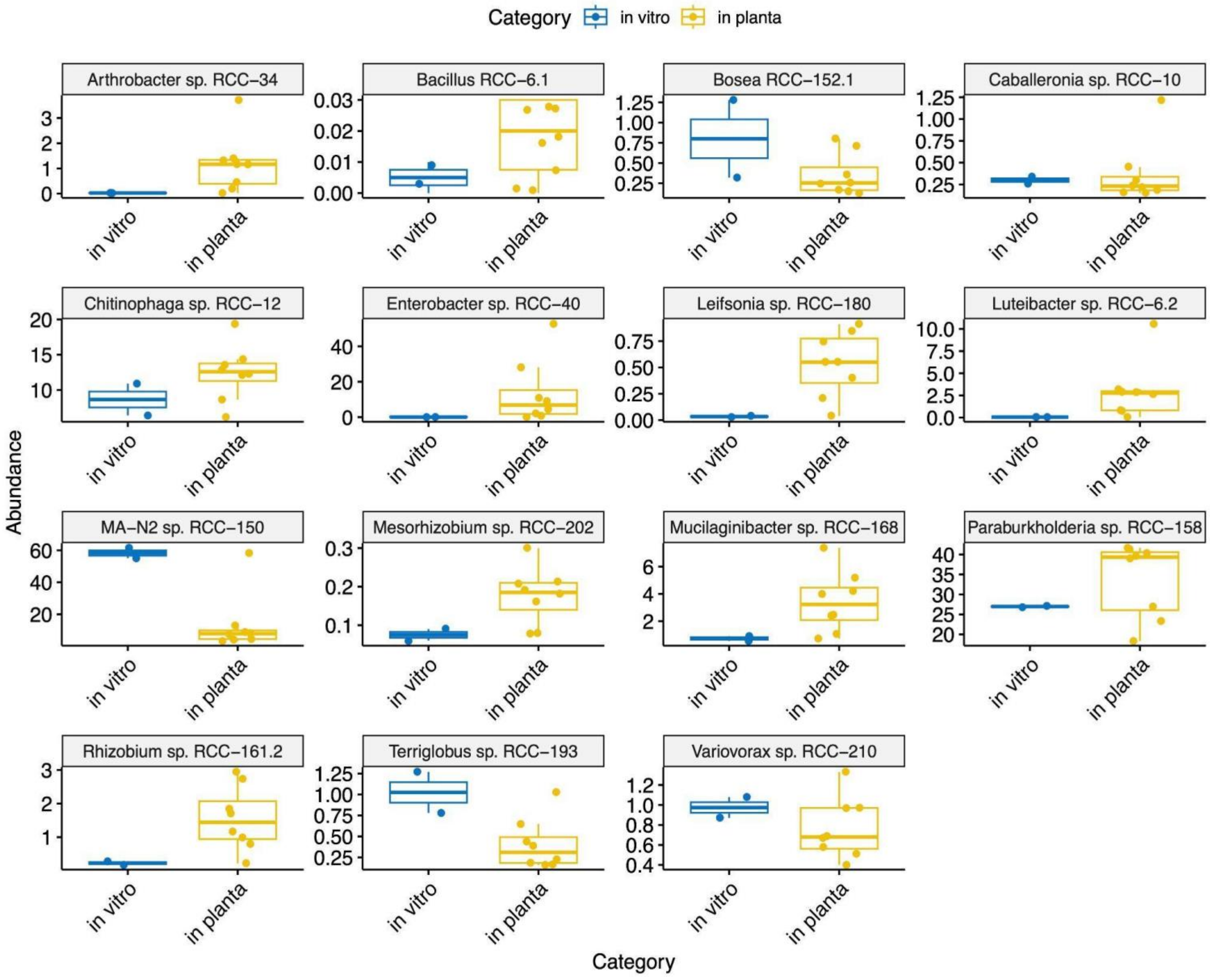


Supplementary Figure S1: Boxplot illustrating the relative abundance and distribution of SynCom members from both *in vitro* and *in planta* experiments, based on a 16S rRNA gene analysis. It highlights the stability in the relative abundance of SynCom members with or without the presence of the rhizosphere. The x-axis represents the experimental treatment, with *in planta* in yellow and *in vitro* in blue, while the y-axis depicts the percentage of relative abundance.

Day 0: Seedlings transferred to the pots

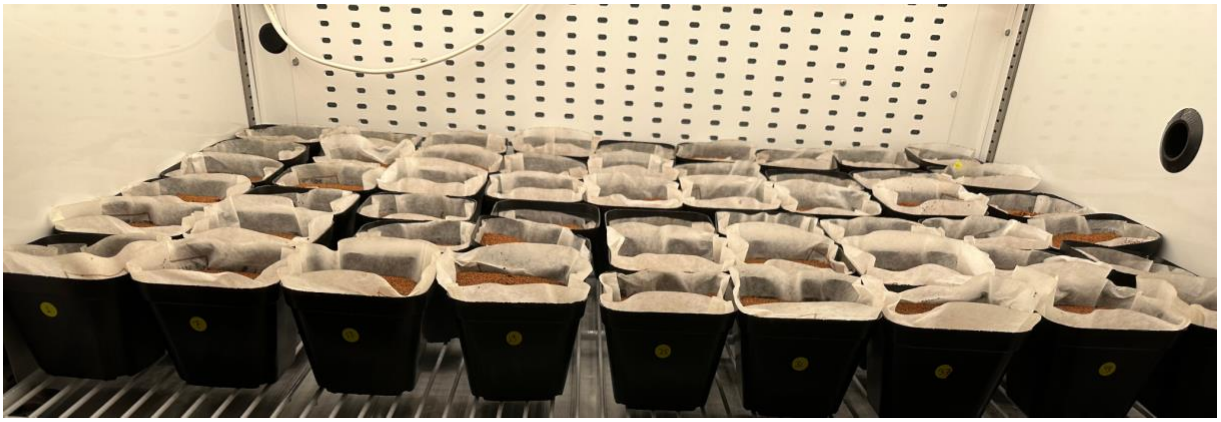

Day7: Seedlings grown in the pot for a week

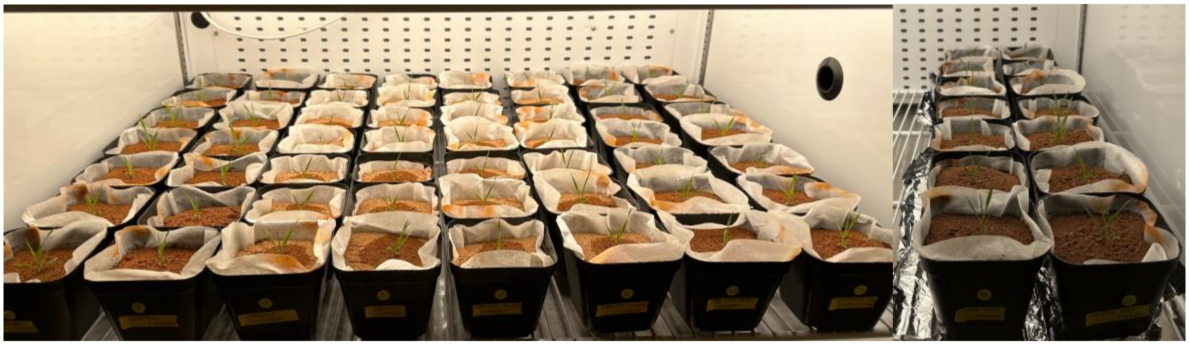

Day 21: Seedlings grown in the pot for 3 weeks before sampling

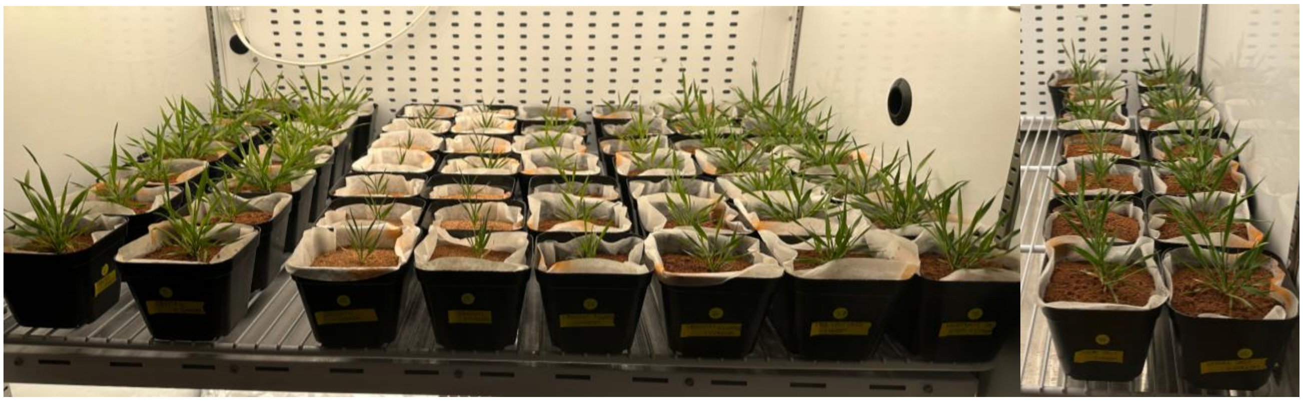

Supplementary Figure S2: Pictures of the pots used in the plant experiment, taken on Day 0, Day 7, and Day 21 (sampling day).

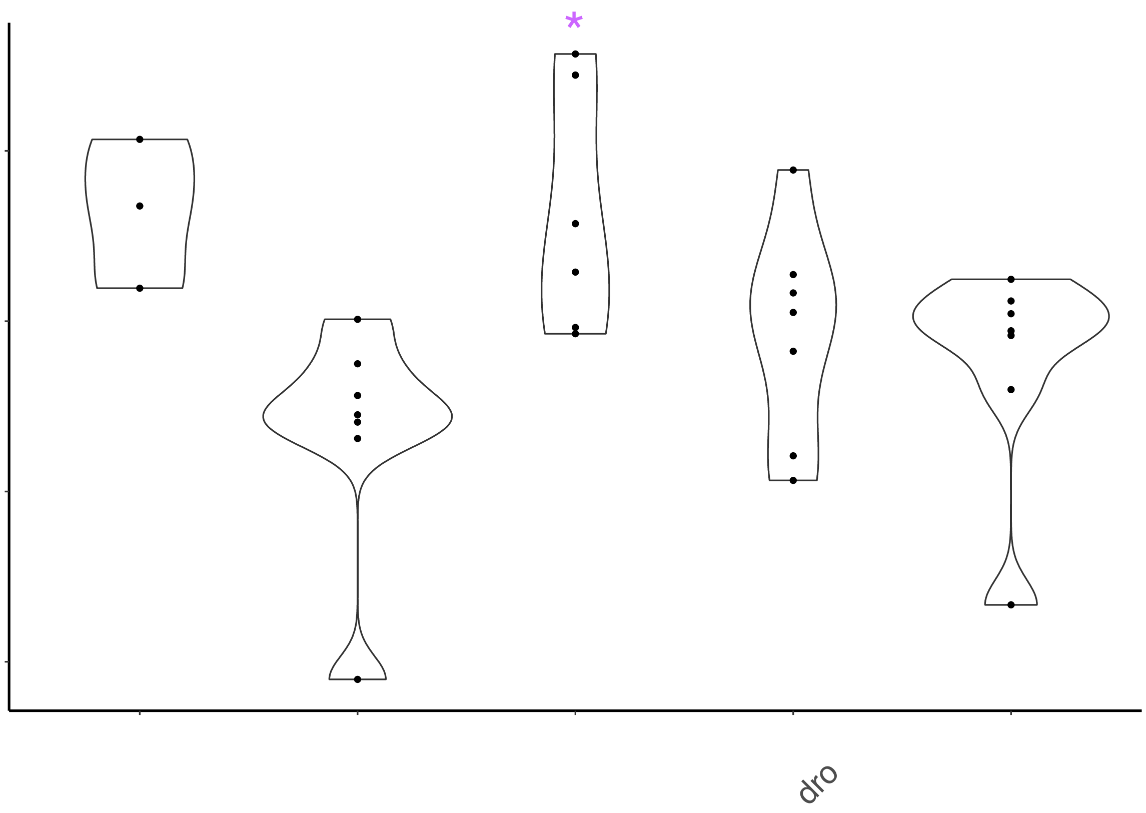


Supplementary Figure S3: Functional redundancy of SynCom isolates calculated using available PGP traits annotated from their genomes. The x-axis represents the inoculum and different *in planta* conditions such as control and stress conditions including drought, rewatered drought, and salt. The y-axis represents the functional redundancy (FR) values estimated using the R package “SYNCSA”. The asterisk indicates significantly different functional redundancy compared to the control.

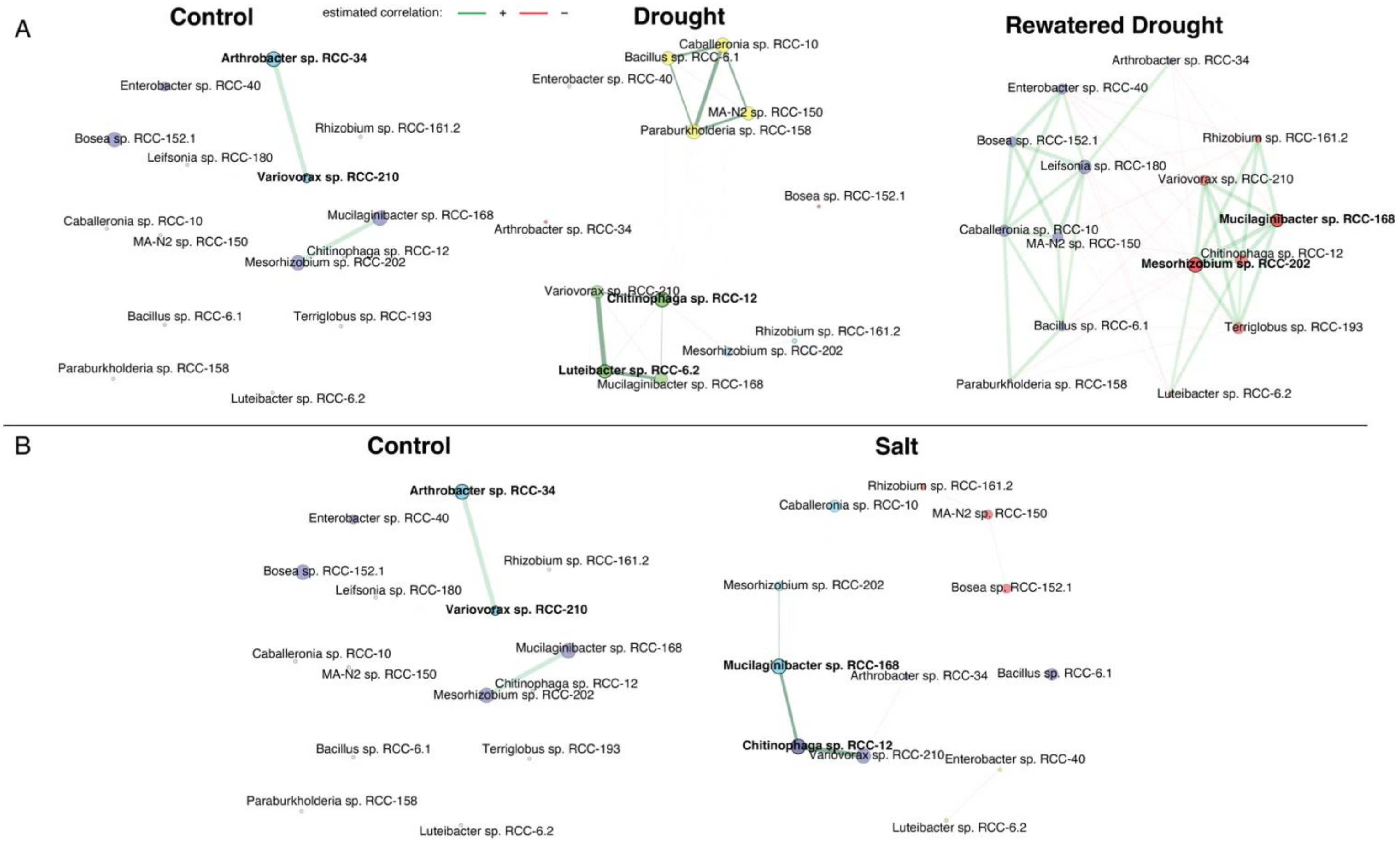


Supplementary Figure S4: Comparison of bacterial association networks between rhizosphere samples under different experimental conditions (A) control versus drought, and (B) control versus salt. R package “NetCoMi” was used to construct and visualize the network. Networks were generated using the fast greedy clustering algorithm with a Pearson correlation coefficient threshold of ± 0.3, and a t-test (< 0.05) for sparse matrix generation. Eigenvector centrality (>0.90) is used for defining hubs and scaling node sizes. Node colors represent clusters and clusters have the same color in both networks if they share at least two taxa. Green edges correspond to positive estimated associations and red edges to negative ones. Hubs are highlighted with bold boundaries and corresponding isolate names.

| Genome | Assembly size (Mbp) | Number of contigs | GC % | Completeness | Contamination | Predicted genes | Phylum | Taxonomy (Based on 16S and produced by SILVA database) | Taxonomy (Based on whole genome and produced by GTDB-tk) |
|---|---|---|---|---|---|---|---|---|---|
| *Caballeronia* sp. RCC-10 | 6.88 | 130 | 62.66 | 99.95 | 0.10 | 6351 | Proteobacteria | Bacteria;Proteobacteria;Gammaproteobacteria;Burkholderiales;Burkholderiaceae;Burkholderia-Caballeronia-Paraburkholderia; | d__Bacteria;p__Proteobacteria;c__Gammaproteobacteria;o__Burkholderiales;f__Burkholderiaceae;g__Caballeronia;s__Caballeronia grimmiae |
| *Chitinophaga* sp. RCC-12 | 8.09 | 34 | 46.82 | 100.00 | 0.49 | 6541 | Bacteroidota | Bacteria;Bacteroidota;Bacteroidia;Chitinophagales;Chitinophagaceae;Chitinophaga; | d__Bacteria;p__Bacteroidota;c__Bacteroidia;o__Chitinophagales;f__Chitinophagaceae;g__Chitinophaga;s__ |
| *MA-N2* sp. RCC-150 | 4.77 | 67 | 63.15 | 99.62 | 2.83 | 4519 | Actinobacteriota | Bacteria;Actinobacteriota;Actinobacteria;Micrococcales;Micrococcaceae; | d__Bacteria;p__Actinobacteriota;c__Actinomycetia;o__Actinomycetales;f__Micrococcaceae;g__MA-N2;s__ |
| *Paraburkholderia* sp. RCC-158 | 8.00 | 95 | 62.62 | 99.95 | 1.59 | 7233 | Proteobacteria | Bacteria;Proteobacteria;Gammaproteobacteria;Burkholderiales;Burkholderiaceae;Burkholderia-Caballeronia-Paraburkholderia; | d__Bacteria;p__Proteobacteria;c__Gammaproteobacteria;o__Burkholderiales;f__Burkholderiaceae;g__Paraburkholderia;s__Paraburkholderia graminis |
| *Rhizobium* sp. RCC-161.2 | 6.91 | 49 | 60.04 | 99.85 | 0.31 | 6585 | Proteobacteria | Bacteria;Proteobacteria;Alphaproteobacteria;Rhizobiales;Rhizobiaceae;Allorhizobium-Neorhizobium-Pararhizobium-Rhizobium; | d__Bacteria;p__Proteobacteria;c__Alphaproteobacteria;o__Rhizobiales;f__Rhizobiaceae;g__Rhizobium;s__ |
| *Mucilaginibacter* sp. RCC-168 | 6.70 | 90 | 42.44 | 97.46 | 0.95 | 5694 | Bacteroidota | Bacteria;Bacteroidota;Bacteroidia;Sphingobacteriales;Sphingobacteriaceae;Mucilaginibacter; | d__Bacteria;p__Bacteroidota;c__Bacteroidia;o__Sphingobacteriales;f__Sphingobacteriaceae;g__Mucilaginibacter;s__ |
| *Leifsonia* sp. RCC-180 | 3.86 | 1 | 69.17 | 99.49 | 0.17 | 3723 | Actinobacteriota | Bacteria;Actinobacteriota;Actinobacteria;Micrococcales;Microbacteriaceae;Leifsonia; | d__Bacteria;p__Actinobacteriota;c__Actinomycetia;o__Actinomycetales;f__Microbacteriaceae;g__Leifsonia;s__ |
| *Terriglobus* sp. RCC-193 | 4.22 | 9 | 57.53 | 100.00 | 1.72 | 3643 | Acidobacteriota | Bacteria; Acidobacteriota; Acidobacteriae; Acidobacteriales; Acidobacteriaceae (Subgroup 1); Terriglobus | d__Bacteria;p__Acidobacteriota;c__Acidobacteriae;o__Acidobacteriales;f__Acidobacteriaceae;g__Terriglobus;s__ |
| *Mesorhizobium* sp. RCC-202 | 6.15 | 29 | 63.30 | 99.92 | 0.41 | 5908 | Proteobacteria | Bacteria;Proteobacteria;Alphaproteobacteria;Rhizobiales;Rhizobiaceae;Mesorhizobium; | d__Bacteria;p__Proteobacteria;c__Alphaproteobacteria;o__Rhizobiales;f__Rhizobiaceae;g__Mesorhizobium;s__ |
| *Variovorax* sp. RCC-210 | 8.52 | 51 | 68.17 | 100.00 | 0.06 | 7730 | Proteobacteria | Bacteria;Proteobacteria;Gammaproteobacteria;Burkholderiales;Comamonadaceae; Variovorax; | d__Bacteria;p__Proteobacteria;c__Gammaproteobacteria;o__Burkholderiales;f__Burkholderiaceae;g__Variovorax;s__Variovorax sp003096925 |
| *Arthrobacter* sp. RCC-34 | 3.67 | 7 | 68.15 | 99.31 | 0.46 | 3343 | Actinobacteriota | Bacteria;Actinobacteriota;Actinobacteria;Micrococcales;Micrococcaceae;Arthrobacter; | d__Bacteria;p__Actinobacteriota;c__Actinomycetia;o__Actinomycetales;f__Micrococcaceae;g__Arthrobacter_E;s__ |
| *Enterobacter* sp. RCC-40 | 4.78 | 18 | 54.69 | 99.89 | 0.08 | 4483 | Proteobacteria | Bacteria;Proteobacteria;Gammaproteobacteria;Enterobacterales;Enterobacteriaceae;Enterobacter; | d__Bacteria;p__Proteobacteria;c__Gammaproteobacteria;o__Enterobacterales;f__Enterobacteriaceae;g__Enterobacter;s__Enterobacter ludwigii |
| *Luteibacter* sp. RCC-6.2 | 4.26 | 35 | 65.61 | 99.66 | 0.39 | 3696 | Proteobacteria | Bacteria;Proteobacteria;Gammaproteobacteria;Xanthomonadales;Rhodanobacteraceae;Luteibacter; | d__Bacteria;p__Proteobacteria;c__Gammaproteobacteria;o__Xanthomonadales;f__Rhodanobacteraceae;g__Luteibacter;s__ |
| *Bacillus* sp. RCC-6.1 | 6.11 | 43 | 34.70 | 99.23 | 0.67 | 6156 | Firmicutes | Bacteria;Firmicutes;Bacilli;Bacillales;Bacillaceae;Bacillus; | d__Bacteria;p__Firmicutes;c__Bacilli;o__Bacillales;f__Bacillaceae_G;g__Bacillus_A;s__Bacillus_A bombysepticus |
| *Bosea* sp. RCC-152.1 | 5.97 | 25 | 66.41 | 99.37 | 0.47 | 5733 | Proteobacteria | Bacteria;Proteobacteria;Alphaproteobacteria;Rhizobiales;Beijerinckiaceae;Bosea; | d__Bacteria;p__Proteobacteria;c__Alphaproteobacteria;o__Rhizobiales;f__Beijerinckiaceae;g__Bosea;s__Bosea vestrisii |

Table S1: General genome features for the 15 SynCom isolates in this study.

| Genome | Betaine synthesis from choline | | Proline catabolism | Trehalose synthesis | Mannitol synthesis |
|---|---|---|---|---|---|
| | Choline dehydrogenase (EC 1.1.99.1) | Betaine-aldehyde dehydrogenase (EC 1.2.1.8) | : dehydrogenase EC:1 | (Presence of at least 3 out of 5 genes) See Table 2 | mannitol-1-phosphate 5-dehydrogenase [EC:1.1.1.17] |
| *Variovorax* sp. RCC-210 | | | | Yes | |
| *Terriglobus* sp. RCC-193 | | | Yes | Yes | |
| *Rhizobium* sp. RCC-161.2 | Yes | Yes | | Yes | |
| *Paraburkholderia* sp. RCC-1 | Yes | Yes | | Yes | |
| *Mucilaginibacter* sp. RCC-168 | | | Yes | Yes | |
| *Mesorhizobium* sp. RCC-202 | Yes | Yes | | | |
| *MA-N2* sp. RCC-150 | Yes | Yes | | Yes | Yes |
| *Luteibacter* sp. RCC-6.2 | | | | Yes | |
| *Leifsonia* sp. RCC-180 | | | | Yes | Yes |
| *Enterobacter* sp. RCC-40 | Yes | Yes | | Yes | Yes |
| *Chitinophaga* sp. RCC-12 | | | Yes | Yes | |
| *Caballeronia* sp. RCC-10 | Yes | Yes | | Yes | |
| *Bosea* RCC-152.1 | | | | Yes | |
| *Bacillus* RCC-6.1 | | | Yes | | |
| *Arthrobacter* sp. RCC-34 | Yes | Yes | | Yes | |

**Table S2: Genes essential for the production of specific osmoprotectants in SynCom isolates.**

| **Genomes** | **Kegg_hit** | **Pfam_hits** |
| --- | --- | --- |
| *Caballeronia* sp. RCC-10 | bile acid:Na+ symporter, BASS family | Sodium Bile acid symporter family [PF01758.19]; SBF-like CPA transporter family (DUF4137) [PF13593.9] |
| *Caballeronia* sp. RCC-10 | F-type H+/Na+-transporting ATPase subunit alpha [EC:7.1.2.2 7.2.2.1] | ATP synthase alpha/beta family, nucleotide-binding domain [PF00006.28]; ATP synthase alpha/beta chain, C terminal domain [PF00306.30]; ATP synthase alpha/beta family, beta-barrel domain [PF02874.26]; Photosystem I psaA/psaB protein [PF00223.22]; ATP synthase [PF00231.22] |
| *Caballeronia* sp. RCC-10 | F-type H+/Na+-transporting ATPase subunit beta [EC:7.1.2.2 7.2.2.1] | ATP synthase alpha/beta family, nucleotide-binding domain [PF00006.28]; ATP synthase alpha/beta family, beta-barrel domain [PF02874.26]; T3SS EscN ATPase C-terminal domain [PF18269.4] |
| *Caballeronia* sp. RCC-10 | putative flavoprotein involved in K+ transport | Flavin-binding monooxygenase-like [PF00743.22]; L-lysine 6-monooxygenase/L-ornithine 5-monooxygenase [PF13434.9]; Pyridine nucleotide-disulphide oxidoreductase [PF07992.17]; FAD-NAD(P)-binding [PF13454.9] |
| *Caballeronia* sp. RCC-10 | potassium-transporting ATPase KdpC subunit | K+-transporting ATPase, c chain [PF02669.18] |
| *Caballeronia* sp. RCC-10 | KUP system potassium uptake protein | K+ potassium transporter [PF02705.19] |
| *Chitinophaga* sp. RCC-12 | | Na+ dependent nucleoside transporter C-terminus [PF07662.16]; Na+ dependent nucleoside transporter N-terminus [PF01773.23] |
| *Chitinophaga* sp. RCC-12 | F-type H+/Na+-transporting ATPase subunit alpha [EC:7.1.2.2 7.2.2.1] | ATP synthase alpha/beta family, nucleotide-binding domain [PF00006.28]; ATP synthase alpha/beta chain, C terminal domain [PF00306.30]; ATP synthase alpha/beta family, beta-barrel domain [PF02874.26]; Photosystem I psaA/psaB protein [PF00223.22] |
| *Chitinophaga* sp. RCC-12 | F-type H+/Na+-transporting ATPase subunit beta [EC:7.1.2.2 7.2.2.1] | ATP synthase alpha/beta family, nucleotide-binding domain [PF00006.28]; ATP synthase alpha/beta family, beta-barrel domain [PF02874.26]; T3SS EscN ATPase C-terminal domain [PF18269.4] |
| *Chitinophaga* sp. RCC-12 | potassium-transporting ATPase KdpC subunit | K+-transporting ATPase, c chain [PF02669.18] |
| *Chitinophaga* sp. RCC-12 | KUP system potassium uptake protein | K+ potassium transporter [PF02705.19] |
| *Chitinophaga* sp. RCC-12 | putative flavoprotein involved in K+ transport | Flavin-binding monooxygenase-like [PF00743.22]; Pyridine nucleotide-disulphide oxidoreductase [PF07992.17]; L-lysine 6-monooxygenase/L-ornithine 5-monooxygenase [PF13434.9]; Lycopene cyclase protein [PF05834.15]; NAD(P)-binding Rossmann-like domain [PF13450.9] |
| *MA-N2* sp. RCC-150 | multicomponent Na+:H+ antiporter subunit D | Proton-conducting membrane transporter [PF00361.23] |
| *MA-N2* sp. RCC-150 | multicomponent Na+:H+ antiporter subunit A | Proton-conducting membrane transporter [PF00361.23]; Domain related to MnhB subunit of Na+/H+ antiporter [PF04039.16]; MBH, subunit E [PF20501.1]; MBH, subunit D [PF13244.9]; NADH-Ubiquinone oxidoreductase (complex I), chain 5 N-terminus [PF00662.23] |
| *MA-N2* sp. RCC-150 | F-type H+/Na+-transporting ATPase subunit beta [EC:7.1.2.2 7.2.2.1] | ATP synthase alpha/beta family, nucleotide-binding domain [PF00006.28]; ATP synthase alpha/beta family, beta-barrel domain [PF02874.26] |

| | | |
|---|---|---|
| *MA-N2* sp. RCC-150 | F-type H+/Na+-transporting ATPase subunit alpha [EC:7.1.2.2 7.2.2.1] | ATP synthase alpha/beta family, nucleotide-binding domain [PF00006.28]; ATP synthase alpha/beta chain, C terminal domain [PF00306.30]; ATP synthase alpha/beta family, beta-barrel domain [PF02874.26]; Photosystem I psaA/psaB protein [PF00223.22] |
| *MA-N2* sp. RCC-150 | putative flavoprotein involved in K+ transport | Flavin-binding monooxygenase-like [PF00743.22]; L-lysine 6-monooxygenase/L-ornithine 5-monooxygenase [PF13434.9]; Pyridine nucleotide-disulphide oxidoreductase [PF07992.17]; Lycopene cyclase protein [PF05834.15]; HI0933-like protein [PF03486.17]; NAD(P)-binding Rossmann-like domain [PF13450.9] |
| *Bosea* RCC-152.1 | phosphate:Na+ symporter | Na+/Pi-cotransporter [PF02690.18]; PhoU domain [PF01895.22] |
| *Bosea* RCC-152.1 | F-type H+/Na+-transporting ATPase subunit alpha [EC:7.1.2.2 7.2.2.1] | ATP synthase alpha/beta family, nucleotide-binding domain [PF00006.28]; ATP synthase alpha/beta chain, C terminal domain [PF00306.30]; ATP synthase alpha/beta family, beta-barrel domain [PF02874.26]; Photosystem I psaA/psaB protein [PF00223.22] |
| *Bosea* RCC-152.1 | putative flavoprotein involved in K+ transport | Flavin-binding monooxygenase-like [PF00743.22]; L-lysine 6-monooxygenase/L-ornithine 5-monooxygenase [PF13434.9]; Pyridine nucleotide-disulphide oxidoreductase [PF07992.17]; HI0933-like protein [PF03486.17] |
| *Bosea* RCC-152.1 | KUP system potassium uptake protein | K+ potassium transporter [PF02705.19] |
| *Bosea* RCC-152.1 | potassium-transporting ATPase KdpC subunit | K+-transporting ATPase, c chain [PF02669.18] |
| *Bosea* RCC-152.1 | potassium-transporting ATPase KdpF subunit | F subunit of K+-transporting ATPase (Potass_KdpF) [PF09604.13] |
| *Paraburkholderia* sp. RCC-158 | phosphate:Na+ symporter | Na+/Pi-cotransporter [PF02690.18] |
| *Paraburkholderia* sp. RCC-158 | concentrative nucleoside transporter, CNT family | Na+ dependent nucleoside transporter C-terminus [PF07662.16]; Na+ dependent nucleoside transporter N-terminus [PF01773.23] |
| *Paraburkholderia* sp. RCC-158 | bile acid:Na+ symporter, BASS family | Sodium Bile acid symporter family [PF01758.19]; SBF-like CPA transporter family (DUF4137) [PF13593.9] |
| *Paraburkholderia* sp. RCC-158 | malate:Na+ symporter | 2-hydroxycarboxylate transporter family [PF03390.18] |
| *Paraburkholderia* sp. RCC-158 | F-type H+/Na+-transporting ATPase subunit beta [EC:7.1.2.2 7.2.2.1] | ATP synthase alpha/beta family, nucleotide-binding domain [PF00006.28]; ATP synthase alpha/beta family, beta-barrel domain [PF02874.26]; T3SS EscN ATPase C-terminal domain [PF18269.4] |
| *Paraburkholderia* sp. RCC-158 | F-type H+/Na+-transporting ATPase subunit alpha [EC:7.1.2.2 7.2.2.1] | ATP synthase alpha/beta family, nucleotide-binding domain [PF00006.28]; ATP synthase alpha/beta chain, C terminal domain [PF00306.30]; ATP synthase alpha/beta family, beta-barrel domain [PF02874.26]; Photosystem I psaA/psaB protein [PF00223.22]; ATP synthase [PF00231.22] |
| *Paraburkholderia* sp. RCC-158 | KUP system potassium uptake protein | K+ potassium transporter [PF02705.19] |
| *Paraburkholderia* sp. RCC-158 | KUP system potassium uptake protein | K+ potassium transporter [PF02705.19] |
| *Paraburkholderia* sp. RCC-158 | putative flavoprotein involved in K+ transport | Flavin-binding monooxygenase-like [PF00743.22]; Pyridine nucleotide-disulphide oxidoreductase [PF07992.17]; L-lysine |

| | | |
|---|---|---|
| | | 6-monooxygenase/L-ornithine 5-monooxygenase [PF13434.9] |
| *Paraburkholderia* sp. RCC-158 | putative flavoprotein involved in K+ transport | Flavin-binding monooxygenase-like [PF00743.22]; L-lysine 6-monooxygenase/L-ornithine 5-monooxygenase [PF13434.9]; Pyridine nucleotide-disulphide oxidoreductase [PF07992.17] |
| *Paraburkholderia* sp. RCC-158 | potassium-transporting ATPase KdpC subunit | K+-transporting ATPase, c chain [PF02669.18] |
| *Rhizobium* sp. RCC-161.2 | F-type H+/Na+-transporting ATPase subunit alpha [EC:7.1.2.2 7.2.2.1] | ATP synthase alpha/beta family, nucleotide-binding domain [PF00006.28]; ATP synthase alpha/beta chain, C terminal domain [PF00306.30]; ATP synthase alpha/beta family, beta-barrel domain [PF02874.26]; Photosystem I psaA/psaB protein [PF00223.22] |
| *Rhizobium* sp. RCC-161.2 | phosphate:Na+ symporter | Na+/Pi-cotransporter [PF02690.18]; PhoU domain [PF01895.22] |
| *Rhizobium* sp. RCC-161.2 | multicomponent Na+:H+ antiporter subunit A | Proton-conducting membrane transporter [PF00361.23]; MBH, subunit E [PF20501.1]; MBH, subunit D [PF13244.9]; NADH-Ubiquinone oxidoreductase (complex I), chain 5 N-terminus [PF00662.23] |
| *Rhizobium* sp. RCC-161.2 | multicomponent Na+:H+ antiporter subunit D | Proton-conducting membrane transporter [PF00361.23] |
| *Rhizobium* sp. RCC-161.2 | potassium-transporting ATPase KdpC subunit | K+-transporting ATPase, c chain [PF02669.18] |
| *Rhizobium* sp. RCC-161.2 | potassium-transporting ATPase KdpF subunit | F subunit of K+-transporting ATPase (Potass_KdpF) [PF09604.13] |
| *Rhizobium* sp. RCC-161.2 | putative flavoprotein involved in K+ transport | Flavin-binding monooxygenase-like [PF00743.22]; Pyridine nucleotide-disulphide oxidoreductase [PF07992.17]; L-lysine 6-monooxygenase/L-ornithine 5-monooxygenase [PF13434.9] |
| *Rhizobium* sp. RCC-161.2 | KUP system potassium uptake protein | K+ potassium transporter [PF02705.19] |
| *Mucilaginibacter* sp. RCC-168 | F-type H+/Na+-transporting ATPase subunit beta [EC:7.1.2.2 7.2.2.1] | ATP synthase alpha/beta family, nucleotide-binding domain [PF00006.28]; ATP synthase alpha/beta family, beta-barrel domain [PF02874.26]; T3SS EscN ATPase C-terminal domain [PF18269.4] |
| *Mucilaginibacter* sp. RCC-168 | bile acid:Na+ symporter, BASS family | Sodium Bile acid symporter family [PF01758.19]; SBF-like CPA transporter family (DUF4137) [PF13593.9] |
| *Mucilaginibacter* sp. RCC-168 | F-type H+/Na+-transporting ATPase subunit alpha [EC:7.1.2.2 7.2.2.1] | ATP synthase alpha/beta family, nucleotide-binding domain [PF00006.28]; ATP synthase alpha/beta chain, C terminal domain [PF00306.30]; ATP synthase alpha/beta family, beta-barrel domain [PF02874.26]; Photosystem I psaA/psaB protein [PF00223.22] |
| *Mucilaginibacter* sp. RCC-168 | potassium-transporting ATPase KdpC subunit | K+-transporting ATPase, c chain [PF02669.18] |
| *Mucilaginibacter* sp. RCC-168 | KUP system potassium uptake protein | K+ potassium transporter [PF02705.19] |
| *Mucilaginibacter* sp. RCC-168 | KUP system potassium uptake protein | K+ potassium transporter [PF02705.19] |
| *Leifsonia* sp. RCC-180 | F-type H+/Na+-transporting ATPase subunit alpha [EC:7.1.2.2 7.2.2.1] | ATP synthase alpha/beta family, nucleotide-binding domain [PF00006.28]; ATP synthase alpha/beta chain, C terminal domain [PF00306.30]; ATP synthase alpha/beta family, beta- |

| | | |
|---|---|---|
| | | barrel domain [PF02874.26]; Photosystem I psaA/psaB protein [PF00223.22] |
| *Leifsonia* sp. RCC-180 | F-type H+/Na+-transporting ATPase subunit beta [EC:7.1.2.2 7.2.2.1] | ATP synthase alpha/beta family, nucleotide-binding domain [PF00006.28]; ATP synthase alpha/beta family, beta-barrel domain [PF02874.26] |
| *Leifsonia* sp. RCC-180 | potassium-transporting ATPase KdpC subunit | K+-transporting ATPase, c chain [PF02669.18] |
| *Leifsonia* sp. RCC-180 | KUP system potassium uptake protein | K+ potassium transporter [PF02705.19] |
| *Leifsonia* sp. RCC-180 | putative flavoprotein involved in K+ transport | Flavin-binding monooxygenase-like [PF00743.22]; Pyridine nucleotide-disulphide oxidoreductase [PF07992.17]; L-lysine 6-monooxygenase/L-ornithine 5-monooxygenase [PF13434.9]; FAD binding domain [PF01494.22]; NAD(P)-binding Rossmann-like domain [PF13450.9]; HI0933-like protein [PF03486.17]; FAD-NAD(P)-binding [PF13454.9]; Lycopene cyclase protein [PF05834.15] |
| *Terriglobus* sp. RCC-193 | F-type H+/Na+-transporting ATPase subunit alpha [EC:7.1.2.2 7.2.2.1] | ATP synthase alpha/beta family, nucleotide-binding domain [PF00006.28]; ATP synthase alpha/beta chain, C terminal domain [PF00306.30]; ATP synthase alpha/beta family, beta-barrel domain [PF02874.26]; Photosystem I psaA/psaB protein [PF00223.22] |
| *Terriglobus* sp. RCC-193 | F-type H+/Na+-transporting ATPase subunit beta [EC:7.1.2.2 7.2.2.1] | ATP synthase alpha/beta family, nucleotide-binding domain [PF00006.28]; ATP synthase alpha/beta family, beta-barrel domain [PF02874.26]; T3SS EscN ATPase C-terminal domain [PF18269.4] |
| *Terriglobus* sp. RCC-193 | concentrative nucleoside transporter, CNT family | Na+ dependent nucleoside transporter C-terminus [PF07662.16]; Na+ dependent nucleoside transporter N-terminus [PF01773.23] |
| *Terriglobus* sp. RCC-193 | potassium-transporting ATPase KdpC subunit | K+-transporting ATPase, c chain [PF02669.18] |
| *Mesorhizobium* sp. RCC-202 | F-type H+/Na+-transporting ATPase subunit alpha [EC:7.1.2.2 7.2.2.1] | ATP synthase alpha/beta family, nucleotide-binding domain [PF00006.28]; ATP synthase alpha/beta chain, C terminal domain [PF00306.30]; ATP synthase alpha/beta family, beta-barrel domain [PF02874.26]; Photosystem I psaA/psaB protein [PF00223.22] |
| *Mesorhizobium* sp. RCC-202 | phosphate:Na+ symporter | Na+/Pi-cotransporter [PF02690.18] |
| *Mesorhizobium* sp. RCC-202 | potassium-transporting ATPase KdpC subunit | K+-transporting ATPase, c chain [PF02669.18] |
| *Mesorhizobium* sp. RCC-202 | potassium-transporting ATPase KdpF subunit | F subunit of K+-transporting ATPase (Potass_KdpF) [PF09604.13] |
| *Mesorhizobium* sp. RCC-202 | putative flavoprotein involved in K+ transport | Flavin-binding monooxygenase-like [PF00743.22]; L-lysine 6-monooxygenase/L-ornithine 5-monooxygenase [PF13434.9]; Pyridine nucleotide-disulphide oxidoreductase [PF07992.17] |
| *Mesorhizobium* sp. RCC-202 | putative flavoprotein involved in K+ transport | Flavin-binding monooxygenase-like [PF00743.22]; L-lysine 6-monooxygenase/L-ornithine 5-monooxygenase [PF13434.9]; Pyridine nucleotide-disulphide oxidoreductase [PF07992.17]; FAD-NAD(P)-binding [PF13454.9] |
| *Mesorhizobium* sp. RCC-202 | putative flavoprotein involved in K+ transport | Flavin-binding monooxygenase-like [PF00743.22]; Pyridine nucleotide-disulphide oxidoreductase [PF07992.17]; L-lysine |

| | | |
|---|---|---|
| | | 6-monooxygenase/L-ornithine 5-monooxygenase [PF13434.9] |
| *Mesorhizobium* sp. RCC-202 | KUP system potassium uptake protein | K+ potassium transporter [PF02705.19] |
| *Mesorhizobium* sp. RCC-202 | KUP system potassium uptake protein | K+ potassium transporter [PF02705.19] |
| *Variovorax* sp. RCC-210 | multicomponent K+:H+ antiporter subunit A | Proton-conducting membrane transporter [PF00361.23]; Domain related to MnhB subunit of Na+/H+ antiporter [PF04039.16]; MBH, subunit E [PF20501.1]; MBH, subunit D [PF13244.9]; NADH-Ubiquinone oxidoreductase (complex I), chain 5 N-terminus [PF00662.23] |
| *Variovorax* sp. RCC-210 | F-type H+/Na+-transporting ATPase subunit beta [EC:7.1.2.2 7.2.2.1] | ATP synthase alpha/beta family, nucleotide-binding domain [PF00006.28]; ATP synthase alpha/beta family, beta-barrel domain [PF02874.26]; T3SS EscN ATPase C-terminal domain [PF18269.4] |
| *Variovorax* sp. RCC-210 | F-type H+/Na+-transporting ATPase subunit alpha [EC:7.1.2.2 7.2.2.1] | ATP synthase alpha/beta family, nucleotide-binding domain [PF00006.28]; ATP synthase alpha/beta chain, C terminal domain [PF00306.30]; ATP synthase alpha/beta family, beta-barrel domain [PF02874.26]; ATP synthase [PF00231.22]; Photosystem I psaA/psaB protein [PF00223.22] |
| *Variovorax* sp. RCC-210 | phosphate:Na+ symporter | Na+/Pi-cotransporter [PF02690.18]; PhoU domain [PF01895.22] |
| *Variovorax* sp. RCC-210 | putative flavoprotein involved in K+ transport | Flavin-binding monooxygenase-like [PF00743.22]; L-lysine 6-monooxygenase/L-ornithine 5-monooxygenase [PF13434.9]; Pyridine nucleotide-disulphide oxidoreductase [PF07992.17]; FAD binding domain [PF01494.22]; Lycopene cyclase protein [PF05834.15] |
| *Variovorax* sp. RCC-210 | multicomponent K+:H+ antiporter subunit D | Proton-conducting membrane transporter [PF00361.23] |
| *Variovorax* sp. RCC-210 | multicomponent K+:H+ antiporter subunit A | Proton-conducting membrane transporter [PF00361.23]; Domain related to MnhB subunit of Na+/H+ antiporter [PF04039.16]; MBH, subunit E [PF20501.1]; MBH, subunit D [PF13244.9]; NADH-Ubiquinone oxidoreductase (complex I), chain 5 N-terminus [PF00662.23] |
| *Variovorax* sp. RCC-210 | KUP system potassium uptake protein | K+ potassium transporter [PF02705.19] |
| *Variovorax* sp. RCC-210 | KUP system potassium uptake protein | K+ potassium transporter [PF02705.19] |
| *Variovorax* sp. RCC-210 | potassium-transporting ATPase KdpC subunit | K+-transporting ATPase, c chain [PF02669.18] |
| *Variovorax* sp. RCC-210 | putative flavoprotein involved in K+ transport | Flavin-binding monooxygenase-like [PF00743.22]; Pyridine nucleotide-disulphide oxidoreductase [PF07992.17] |
| *Arthrobacter* sp. RCC-34 | F-type H+/Na+-transporting ATPase subunit alpha [EC:7.1.2.2 7.2.2.1] | ATP synthase alpha/beta family, nucleotide-binding domain [PF00006.28]; ATP synthase alpha/beta chain, C terminal domain [PF00306.30]; ATP synthase alpha/beta family, beta-barrel domain [PF02874.26]; Photosystem I psaA/psaB protein [PF00223.22] |
| *Arthrobacter* sp. RCC-34 | F-type H+/Na+-transporting ATPase subunit beta [EC:7.1.2.2 7.2.2.1] | ATP synthase alpha/beta family, nucleotide-binding domain [PF00006.28]; ATP synthase alpha/beta family, beta-barrel domain [PF02874.26] |
| *Arthrobacter* sp. RCC-34 | multicomponent Na+:H+ antiporter subunit D | Proton-conducting membrane transporter [PF00361.23] |

| | | |
|---|---|---|
| *Arthrobacter* sp. RCC-34 | multicomponent Na+:H+ antiporter subunit A | Proton-conducting membrane transporter [PF00361.23]; Domain related to MnhB subunit of Na+/H+ antiporter [PF04039.16]; MBH, subunit E [PF20501.1]; MBH, subunit D [PF13244.9]; NADH-Ubiquinone oxidoreductase (complex I), chain 5 N-terminus [PF00662.23] |
| *Arthrobacter* sp. RCC-34 | bile acid:Na+ symporter, BASS family | Sodium Bile acid symporter family [PF01758.19]; SBF-like CPA transporter family (DUF4137) [PF13593.9] |
| *Arthrobacter* sp. RCC-34 | potassium-transporting ATPase KdpC subunit | K+-transporting ATPase, c chain [PF02669.18] |
| *Enterobacter* sp. RCC-40 | nucleoside transport protein | Na+ dependent nucleoside transporter C-terminus [PF07662.16]; Na+ dependent nucleoside transporter N-terminus [PF01773.23] |
| *Enterobacter* sp. RCC-40 | Na+:H+ antiporter, NhaB family | Bacterial Na+/H+ antiporter B (NhaB) [PF06450.15]; Citrate transporter [PF03600.19] |
| *Enterobacter* sp. RCC-40 | nucleoside transport protein | Na+ dependent nucleoside transporter C-terminus [PF07662.16]; Na+ dependent nucleoside transporter N-terminus [PF01773.23] |
| *Enterobacter* sp. RCC-40 | Na+-transporting NADH:ubiquinone oxidoreductase subunit A [EC:7.2.1.1] | Na(+)-translocating NADH-quinone reductase subunit A (NQRA) [PF05896.14]; NQRA C-terminal domain [PF11973.11] |
| *Enterobacter* sp. RCC-40 | Na+-transporting NADH:ubiquinone oxidoreductase subunit B [EC:7.2.1.1] | NQR2, RnfD, RnfE family [PF03116.18] |
| *Enterobacter* sp. RCC-40 | Na+-transporting NADH:ubiquinone oxidoreductase subunit C [EC:7.2.1.1] | FMN-binding domain [PF04205.17] |
| *Enterobacter* sp. RCC-40 | Na+-transporting NADH:ubiquinone oxidoreductase subunit D [EC:7.2.1.1] | Rnf-Nqr subunit, membrane protein [PF02508.17] |
| *Enterobacter* sp. RCC-40 | Na+-transporting NADH:ubiquinone oxidoreductase subunit E [EC:7.2.1.1] | Rnf-Nqr subunit, membrane protein [PF02508.17] |
| *Enterobacter* sp. RCC-40 | Na+-transporting NADH:ubiquinone oxidoreductase subunit F [EC:7.2.1.1] | Oxidoreductase NAD-binding domain [PF00175.24]; Oxidoreductase FAD-binding domain [PF00970.27]; 2Fe-2S iron-sulfur cluster binding domain [PF00111.30] |
| *Enterobacter* sp. RCC-40 | malate:Na+ symporter | 2-hydroxycarboxylate transporter family [PF03390.18] |
| *Enterobacter* sp. RCC-40 | F-type H+/Na+-transporting ATPase subunit alpha [EC:7.1.2.2 7.2.2.1] | ATP synthase alpha/beta family, nucleotide-binding domain [PF00006.28]; ATP synthase alpha/beta chain, C terminal domain [PF00306.30]; ATP synthase alpha/beta family, beta-barrel domain [PF02874.26]; Photosystem I psaA/psaB protein [PF00223.22]; ATP synthase [PF00231.22] |
| *Enterobacter* sp. RCC-40 | F-type H+/Na+-transporting ATPase subunit beta [EC:7.1.2.2 7.2.2.1] | ATP synthase alpha/beta family, nucleotide-binding domain [PF00006.28]; ATP synthase alpha/beta family, beta-barrel domain [PF02874.26]; T3SS EscN ATPase C-terminal domain [PF18269.4] |
| *Enterobacter* sp. RCC-40 | bile acid:Na+ symporter, BASS family | Sodium Bile acid symporter family [PF01758.19]; SBF-like CPA transporter family (DUF4137) [PF13593.9] |
| *Enterobacter* sp. RCC-40 | phosphate:Na+ symporter | Na+/Pi-cotransporter [PF02690.18] |
| *Enterobacter* sp. RCC-40 | potassium-transporting ATPase KdpC subunit | K+-transporting ATPase, c chain [PF02669.18] |
| *Enterobacter* sp. RCC-40 | KUP system potassium uptake protein | K+ potassium transporter [PF02705.19] |
| *Bacillus* RCC-6.1 | phosphate:Na+ symporter | Na+/Pi-cotransporter [PF02690.18]; PhoU domain [PF01895.22] |

| | | |
|---|---|---|
| *Bacillus* RCC-6.1 | nucleoside transport protein | Na+ dependent nucleoside transporter C-terminus [PF07662.16]; Na+ dependent nucleoside transporter N-terminus [PF01773.23] |
| *Bacillus* RCC-6.1 | nucleoside transport protein | Na+ dependent nucleoside transporter C-terminus [PF07662.16]; Na+ dependent nucleoside transporter N-terminus [PF01773.23] |
| *Bacillus* RCC-6.1 | malate:Na+ symporter | 2-hydroxycarboxylate transporter family [PF03390.18] |
| *Bacillus* RCC-6.1 | putative amino acid transporter | Na+-H+ antiporter family [PF13726.9]; Na+/H+ antiporter family [PF03553.17] |
| *Bacillus* RCC-6.1 | nucleoside transport protein | Na+ dependent nucleoside transporter C-terminus [PF07662.16]; Na+ dependent nucleoside transporter N-terminus [PF01773.23] |
| *Bacillus* RCC-6.1 | nucleoside transport protein | Na+ dependent nucleoside transporter C-terminus [PF07662.16]; Na+ dependent nucleoside transporter N-terminus [PF01773.23] |
| *Bacillus* RCC-6.1 | concentrative nucleoside transporter, CNT family | Na+ dependent nucleoside transporter C-terminus [PF07662.16]; Na+ dependent nucleoside transporter N-terminus [PF01773.23] |
| *Bacillus* RCC-6.1 | F-type H+/Na+-transporting ATPase subunit beta [EC:7.1.2.2 7.2.2.1] | ATP synthase alpha/beta family, nucleotide-binding domain [PF00006.28]; ATP synthase alpha/beta family, beta-barrel domain [PF02874.26]; T3SS EscN ATPase C-terminal domain [PF18269.4] |
| *Bacillus* RCC-6.1 | F-type H+/Na+-transporting ATPase subunit alpha [EC:7.1.2.2 7.2.2.1] | ATP synthase alpha/beta family, nucleotide-binding domain [PF00006.28]; ATP synthase alpha/beta chain, C terminal domain [PF00306.30]; ATP synthase alpha/beta family, beta-barrel domain [PF02874.26]; Photosystem I psaA/psaB protein [PF00223.22] |
| *Bacillus* RCC-6.1 | bile acid:Na+ symporter, BASS family | Sodium Bile acid symporter family [PF01758.19]; SBF-like CPA transporter family (DUF4137) [PF13593.9] |
| *Bacillus* RCC-6.1 | purine nucleoside transport protein | Na+ dependent nucleoside transporter C-terminus [PF07662.16]; Na+ dependent nucleoside transporter N-terminus [PF01773.23] |
| *Bacillus* RCC-6.1 | | Na+ dependent nucleoside transporter C-terminus [PF07662.16]; Na+ dependent nucleoside transporter N-terminus [PF01773.23] |
| *Bacillus* RCC-6.1 | potassium-transporting ATPase KdpC subunit | K+-transporting ATPase, c chain [PF02669.18] |
| *Bacillus* RCC-6.1 | putative flavoprotein involved in K+ transport | Flavin-binding monooxygenase-like [PF00743.22]; L-lysine 6-monooxygenase/L-ornithine 5-monooxygenase [PF13434.9]; Pyridine nucleotide-disulphide oxidoreductase [PF07992.17]; NAD(P)-binding Rossmann-like domain [PF13450.9]; HI0933-like protein [PF03486.17]; FAD dependent oxidoreductase [PF01266.27] |
| *Luteibacter* sp. RCC-6.2 | concentrative nucleoside transporter, CNT family | Na+ dependent nucleoside transporter C-terminus [PF07662.16]; Na+ dependent nucleoside transporter N-terminus [PF01773.23] |
| *Luteibacter* sp. RCC-6.2 | F-type H+/Na+-transporting ATPase subunit alpha [EC:7.1.2.2 7.2.2.1] | ATP synthase alpha/beta family, nucleotide-binding domain [PF00006.28]; ATP synthase alpha/beta chain, C terminal domain [PF00306.30]; ATP synthase alpha/beta family, beta-barrel domain [PF02874.26]; Photosystem I psaA/psaB protein [PF00223.22] |

| | | |
|---|---|---|
| *Luteibacter* sp. RCC-6.2 | F-type H+/Na+-transporting ATPase subunit beta [EC:7.1.2.2 7.2.2.1] | ATP synthase alpha/beta family, nucleotide-binding domain [PF00006.28]; ATP synthase alpha/beta family, beta-barrel domain [PF02874.26]; T3SS EscN ATPase C-terminal domain [PF18269.4] |
| *Luteibacter* sp. RCC-6.2 | potassium-transporting ATPase KdpC subunit | K+-transporting ATPase, c chain [PF02669.18] |
| *Luteibacter* sp. RCC-6.2 | potassium-transporting ATPase KdpF subunit | F subunit of K+-transporting ATPase (Potass_KdpF) [PF09604.13] |
| *Luteibacter* sp. RCC-6.2 | KUP system potassium uptake protein | K+ potassium transporter [PF02705.19] |

**Table S3: List of Sodium (Na+) and potassium (K+) transporters present in the genomes of SynCom isolates.**
